# Reductive Enzyme Cascades for Valorization of PET Deconstruction Products

**DOI:** 10.1101/2022.12.16.520786

**Authors:** Madan R. Gopal, Roman M. Dickey, Neil D. Butler, Michael R. Talley, Ashlesha Mohapatra, Mary P. Watson, Wilfred Chen, Aditya M. Kunjapur

## Abstract

To better incentivize the collection of plastic wastes, new chemical transformations must be developed that add value to plastic deconstruction products. Polyethylene terephthalate (PET) is a common plastic whose deconstruction through chemical or biological means has received much attention. However, a limited number of alternative products have been formed from PET deconstruction, and only a small share could serve as building blocks for alternative materials or therapeutics. Here, we demonstrate the production of useful mono-amine and diamine building blocks from known PET deconstruction products. We achieve this by designing one-pot biocatalytic transformations that are informed by the substrate specificity of an ω-transaminase and diverse carboxylic acid reductases (CAR) towards PET deconstruction products. We first establish that an ω-transaminase from *Chromobacterium violaceum* (cvTA) can efficiently catalyze amine transfer to potential PET-derived aldehydes to form the mono-amine *para*-(aminomethyl)benzoic acid (pAMBA) or the diamine *para*-xylylenediamine (pXYL). We then identified CAR orthologs that could perform the bifunctional reduction of TPA to terephthalaldehyde (TPAL) or the reduction of *mono*-(2-hydroxyethyl) terephthalic acid (MHET) to its corresponding aldehyde. After characterizing 17 CARs *in vitro*, we show that the CAR from *Segniliparus rotundus* (srCAR) had the highest observed activity on TPA. Given these newly elucidated substrate specificity results, we designed modular enzyme cascades based on coupling srCAR and cvTA in one-pot with enzymatic co-factor regeneration. When we supply TPA, we achieve a 69 ± 1% yield of pXYL, which is useful as a building block for materials. When we instead supply MHET and subsequently perform base-catalyzed ester hydrolysis, we achieve 70 ± 8% yield of pAMBA, which is useful for therapeutic applications and as a pharmaceutical building block. This work expands the breadth of products derived from PET deconstruction and lays the groundwork for eventual valorization of waste PET to higher-value chemicals and materials.

**GRAPHICAL ABSTRACT:** 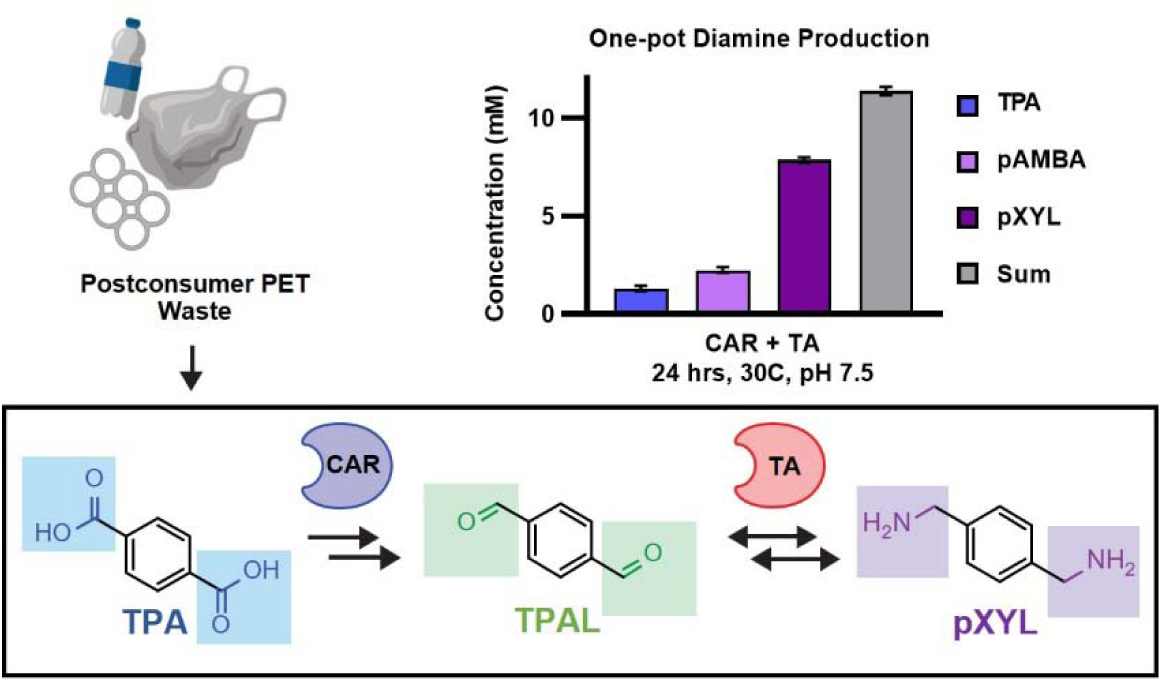

## Introduction

The future of plastics production will require greater circularity to decrease dependence on petroleum as a raw material and to stem the ever-growing flow of plastic waste into landfills.^1–5^ Mechanical recycling of plastics has been the predominant technology to enable the reuse of postconsumer plastic waste. However, chemical deconstruction by synthetic or enzymatic approaches has emerged to allow for recovery of the monomer building blocks of various polymer plastics.^6,7^ While chemical deconstruction offers a simple path to remake the original, virgin-like polymer, broader adoption and utility of chemical deconstruction will benefit from the design of sustainable methods to add value to deconstruction products, making them amenable to upcycled applications.^3,8^ One of the most widely used single-use plastics is polyethylene terephthalate (PET), a member of the thermoplastic category of polymers.^9^ Chemical and enzymatic deconstruction methods have been applied to polyethylene terephthalate (PET) to produce products such as *bis*-(2-hydroxyethyl) terephthalate (BHET), *mono*-(2- hydroxyethyl) terephthalic acid (MHET), and terephthalic acid (TPA).^10–17^ To date, most efforts have harnessed these compounds for either the resynthesis of PET, biological degradation as catabolized carbon sources, or aliphatic building blocks derived from aromatic ring cleavage.^18–20^

We wondered whether we could design a route to convert PET deconstruction products to upcycled amines as alternatives to these products, while preserving the aromatic nature of the terephthalate monomer (**Fig. 1A)**.^21^ Diamines such as *para*-xylylenediamine (pXYL) are value-added monomers for both thermoset and thermoplastic polymers. pXYL can be a component of polyamides, polyimides, or non-isocyanate polyurethanes (**Fig. S1**).^22,23^ Such materials would substantially expand the breadth of products derived from PET deconstruction if an environmentally friendly option were available for valorization. However, chemical synthesis of pXYL requires multiple steps, elevated temperatures and pressures, and strong organic solvents.^24,25^ While diamines can be used as value-added monomers, a significant opportunity exists for the valorization of PET-derived monomers to mono-amines such as *para*-(aminomethyl)benzoic acid (pAMBA), an antifibrinolytic drug used to promote blood clotting and treat fibrotic skin conditions.^26^ Monofunctional molecules are often challenging to make from bifunctional substrates (*e.g.,* TPA), resulting in poor atom economy and low selectivity.^27^ However, enzymatic PET deconstruction offers a previously untapped potential to leverage substrates like MHET, a unique carboxylate with a ester “protecting” group, which could allow for monofunctionalization of terephthalate.

**Figure 1.**
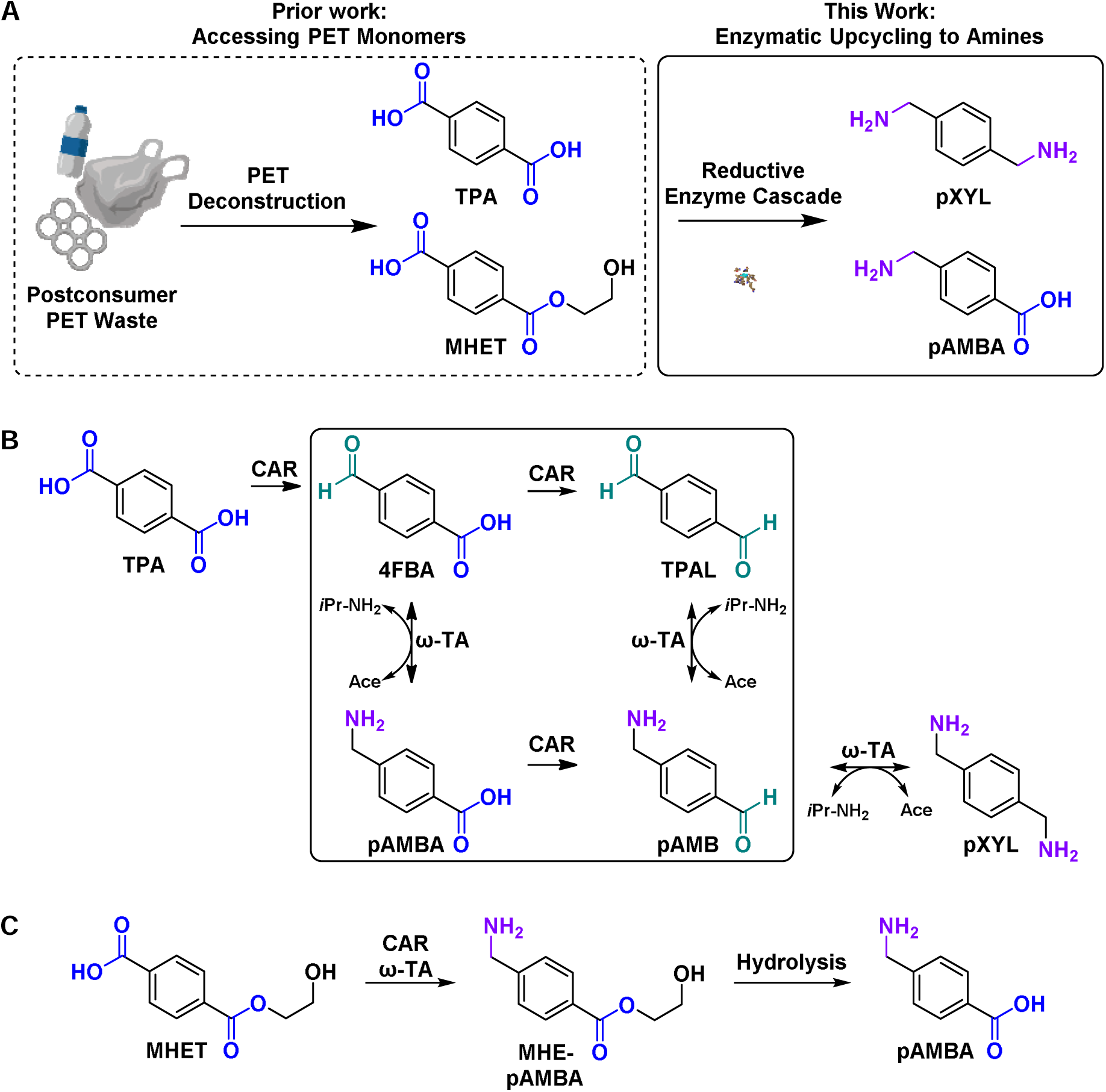
Strategies for valorizing deconstruction products of PET. (A) We envision using this enzyme cascade to upcycle PET waste by creating value-added amines for downstream application in therapeutics and polymer synthesis (B) Envisioned reaction schematic for the one-pot reduction and amination of TPA to pXYL by an enzyme cascade consisting of carboxylic acid reductases (CAR) and ω-transaminases (ω-TA). The reaction cascade goes through four potential intermediate compounds, 4FBA, TPAL, pAMBA, and pAMBA, due to successive reduction and amination reactions. (C) The substrate specificities of CAR and ω-TA could allow functionalization of MHET to pAMBA, a small-molecule therapeutic, by a simple one-pot enzymatic cascade. *i*Pr-NH_2_, isopropylamine; CAR, carboxylic acid reductase; ω-TA, ω-transaminase; Ace, acetone; TPA, terephthalic acid; 4FBA, 4-formylbenzoic acid; TPAL, terephthalaldehyde; pAMBA, *para-*(aminomethyl)benzoic acid; pAMB, *para-*aminomethylbenzaldehyde; pXYL, *para*-xylylenediamine.

Biocatalysis presents an industrially established and green alternative for synthesis of amines and diamines from carbonyl-containing precursors.^28^ Here, we envisioned an enzyme cascade that could convert TPA to pXYL in one pot (**Fig. 1B**), and we wondered whether it could be adjusted such that it could instead generate pAMBA **(Fig. 1C)**. Our retrobiosynthetic design focused on steps of aldehyde consumption and aldehyde generation.^29,30^ Pyridoxal 5’-phosphate- (PLP)-dependent ω-transaminases (ω-TA, EC 2.6.1.x) have been previously applied for the conversion of aldehydes to amines using simple co-substrates like isopropylamine (*i*Pr-NH_2_), though to our knowledge, these enzymes have not been reported to have activity on bifunctional aromatic aldehydes.^31–33^ As PET deconstruction liberates carboxylates rather than aldehydes, we hypothesized that carboxylic acid reductases (CAR, E.C. 1.2.1.30) may be able to catalyze the reduction of carboxylates like TPA to its corresponding dialdehyde, terephthalaldehyde (TPAL).^34–36^ CARs are multidomain and polyspecific enzymes that perform the desirable 2e^-^ reduction of carboxylic acids to aldehydes at a cost of ATP and NADPH.^37^ Given the broad substrate scope of both CAR and ω-TA families, some precedent for this type of cascade exists for certain aliphatic or heterocyclic chemistries.^38,39^ In 2019, Fedorchuk et al. designed a CAR and ω-TA cascade for one-pot transformation of the dicarboxylate adipic acid to hexamethylenediamine, a diamine precursor to nylon.^40^ That study achieved only 30% yield, even after significant engineering to develop two CAR variants for use alongside two distinct transaminase orthologs. A major challenge was that each unique CAR or TA variant could accept a mono-functionalized intermediate or a bifunctionalized intermediate but not both.

Here, we report the design of enzyme cascades that feature single ω-TA and CAR variants to produce amines at high selectivity and yield in a one pot reaction from PET-derived monomers. We examined the specificity of the ω-TA from *Chromobacterium violaceum* (cvTA), which successfully accepted several PET-derived aldehydes while harnessing *i*Pr-NH_2_ as an amine donor. We next explored the diversity of natural CARs using a protein sequence similarity network to identify known and novel putative CARs with activity on TPA. Our study revealed unexpected substrate specificity that enabled us to design highly selective routes to pXYL or to pAMBA by coupling the CAR from *Segniliparus rotundus* (srCAR) to cvTA. Our work represents a notable advance in the green conversion of PET deconstruction products to aromatic (di)aldehydes and (di)amines that could serve as platform intermediates for value-added polymeric materials or pharmaceuticals. By doing so, our work expands the sparse list of biological tools currently available for valorization of plastic deconstruction streams.

## Results and Discussion

### An ω-Transaminase from *Chromobacterium violaceum* (cvTA) Accepts PET-derived Aldehydes

From our retrobiosynthetic analysis, we hypothesized that ω-transaminases could catalyze synthesis of diamines and monoamines from PET-derived aromatic aldehydes. We chose the model ω-TA from *Chromobacterium violaceum* (cvTA) because of its wide substrate scope, well-characterized reaction mechanism, retention of activity in organic cosolvents, and compatibility with the simple amine donor, isopropylamine (*i*Pr-NH_2_).^41–43^ After identifying the reactions in which the ω-TA could participate (**Fig. 2A**), we expressed, purified, and tested cvTA activity on 4-formylbenzoic acid (4FBA) and TPAL using *i*Pr-NH_2_ as a co-substrate. After addition of 10 mM of TPAL to the reaction buffer containing 1.5 μM of cvTA, 40 mM *i*Pr-NH_2_, 5% DMSO, 100 mM HEPES pH 7.5, and 400 μM PLP, we noticed initial turbidity in our reaction. However, we were pleased to see that after incubation for 24 h at 30 °C, the reaction volume became translucent, and HPLC-UV analysis indicated complete conversion of TPAL to the desired diamine pXYL (**Fig. 2B**). Under the same reaction conditions used for pXYL formation, we also observed complete conversion of 10 mM 4FBA to pAMBA (**Fig. 2C**). The highly efficient conversion of candidate aldehyde substrates to amine products catalyzed by cvTA led us to next wonder if CARs could generate these aldehydes from PET deconstruction product.

**Figure 2.**
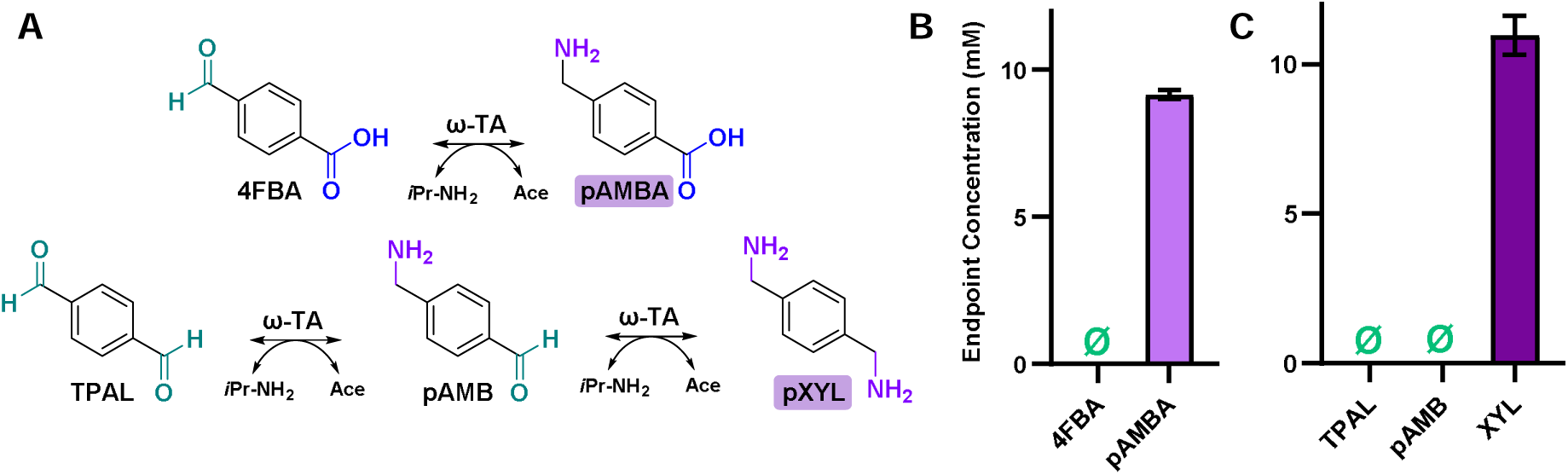
Testing cvTA on PET-derived aldehydes to produce amines. (A) cvTA reactions for the conversion 4FBA and TPAL to their corresponding amines. (B) Endpoint bioconversion of 10 mM 4FBA to pAMBA and (C) 10 mM TPAL to pXYL by 1.5 μM cvTA in 100mM HEPES pH 7.5, 400 μM PLP with 40 mM *i*Pr-NH_2_ as the amine donor. This reaction proceeds to completion after 24 h at 30 °C, with no detection of aldehyde reactants or intermediates after 24 h. Sample size is n=3 and all data are shown as the mean ± standard deviation.

### Bioprospecting Known and Putative CARs for Activity on Potential PET Deconstruction Products

While CARs are extremely useful enzymes, bioprospecting for CAR remains relatively underexplored, likely due to many factors including only recent elucidation of its structure and catalytic cycle, cost of synthesis and assembly of the >3500 bp CAR gene, and the need for stoichiometric equivalents of NADPH and ATP per catalytic cycle.^37,44,45^ To address this gap in the literature, and to identify CARs with specificity and activity towards products of PET deconstruction, we sought to investigate the natural CAR landscape. We first created a protein sequence similarity network (SSN) using the Enzyme Function Initiative Enzyme Similarity Tool^46^ and populated the SSN by selecting five previously characterized CARs, with at least one from each of four phylogenetic types of CARs:^47^ (i) Type I CAR from *Nocardia iowensis* (niCAR);^48^ (ii) Type II CAR from *Stachybotrys bisby* (sbCAR);^49^ (iii) Type III CARs from *Neurospora crassa* (ncCAR) and *Thermothelomyces thermophila* (ttCAR);^50,51^ (iv) Type IV CAR from *Trametes versicolor* (tvCAR2).^52^

For each of these CARs, we performed standard protein BLAST against the NCBI non-redundant protein sequences database. We ultimately populated the SSN with 500 sequences and visualized it using Cytoscape,^53^ where we observed clear distinction between bacterial CARs and fungal type II, III, and IV CARs. The SSN also shows clusters of related enzymes classes known as L-aminoadipate semialdehyde dehydrogenases (E.C. 1.2.1.31)^54^ and L-aminoadipate reductases (E.C. 1.2.1.95) (**Fig. 3A**).^55^ Based on this representative network, we selected 22 putative CARs for cloning and further investigation: 10 from the cluster occupied by the promiscuous Type I bacterial CARs (five previously uncharacterized); three from the presumed Type II cluster (two previously uncharacterized); four from the presumed Type III cluster (two previously uncharacterized); one putative enzyme from *Aspergillus fumigatus* from the less closely related portion of the Type III CAR cluster; one putative enzyme from the presumed Type IV cluster; and three non-clustered, uncharacterized putative enzymes from distinct eukaryotes (**Table S1**).

**Figure 3.**
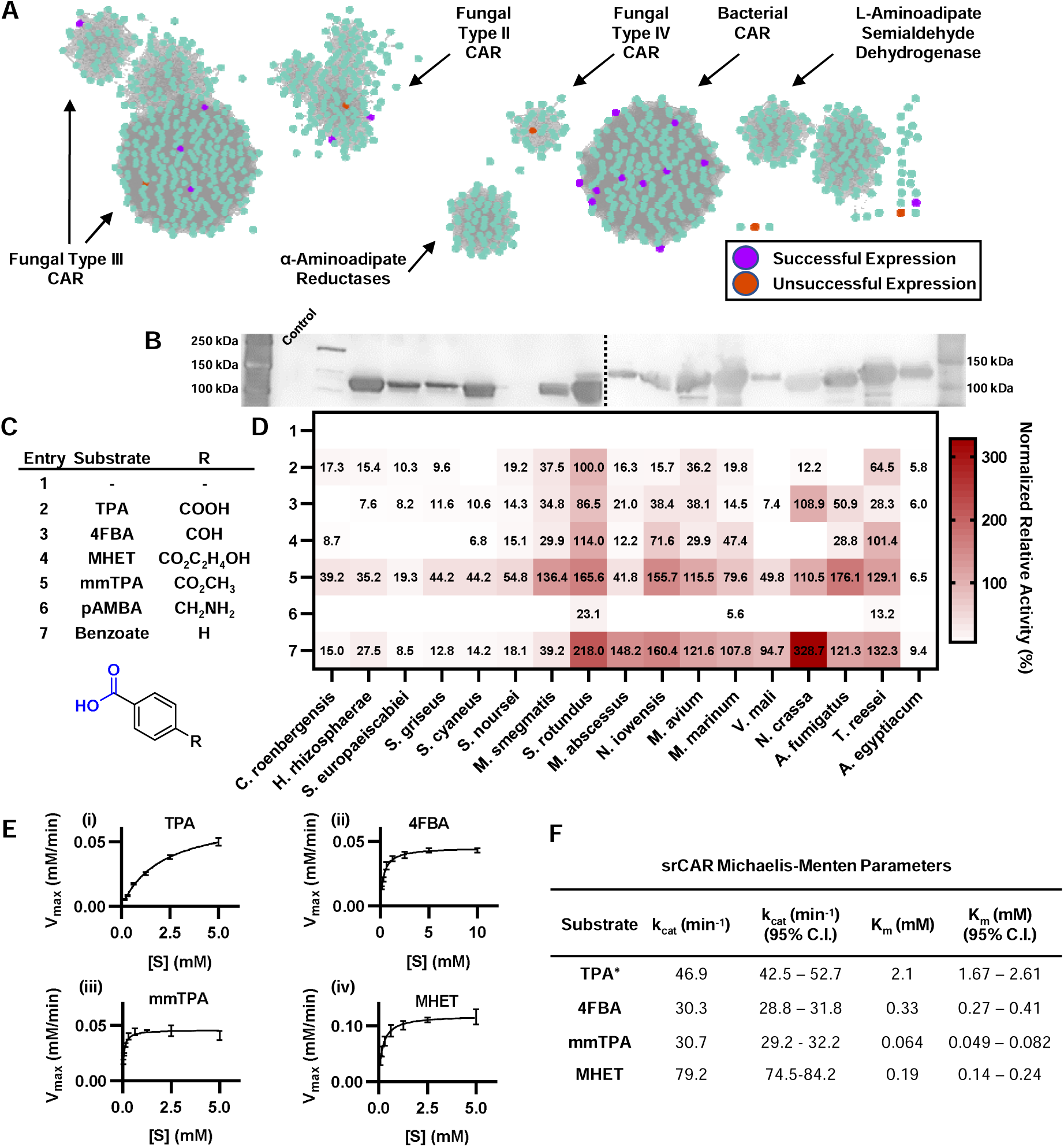
Evaluating CAR specificity on products of PET deconstruction. (A) Using a sequence similarity network, we sampled CARs across distinct protein clusters to probe for activity on TPA. Of 22 cloned CARs, 17 expressed, with 9 of those CARs being unreported in literature. (B) Anti-His_6_ Western blot of cell lysate from *E. coli* BL21 (DE3) harboring different CARs is shown by phylogeny. CARs were expressed under optimized conditions. The Western blot is comprised of two separate blots, which are delineated by the dashed lines and contain their own protein ladders. The first lane of the Western blot is a control lane consisting of lysate from *E. coli* BL21 (DE3) with no CAR plasmid. A band corresponding to the putative CAR from *Streptomyces noursei* was not detected during Western blot analysis but was confirmed in a separate experiment. (C) We evaluated the substrate specificity of CAR on TPA, 4FBA, MHET and mmTPA, which are carboxylates related to PET depolymerization. In addition, we tested pAMBA, as it is a potential reaction intermediate in our proposed multienzyme cascade for the formation of amines. (D) Under mild aqueous conditions, we tracked the conversion of CAR on TPA, 4FBA, MHET, mmTPA, pAMBA and the model aromatic acid, benzoic acid. Here, the oxidation of NADPH is measured as a proxy to aldehyde formation, as depletion of NADPH is linked to the consumption of substrate. The heatmap was generated by calculating the triplicate average of initial rates of each CAR-substrate pair. The heatmap values represent the average initial rate, normalized to the average activity of srCAR on TPA of 4.80 mM NADPH_ox_·h^-1^·μM ^-1^. CARs-substrate pairs with <5% normalized relative activity are not shown (E) A Michaelis-Menten study was performed on the highest performing CAR from *Segniliparus rotundus* to evaluate k_cat_ and K_m_ on TPA (i), 4FBA (ii), MHET (iii) and mmTPA (iv). (F) Michaelis-Menten parameters and 95% confidence intervals were fitted using non-linear least squares regression on Graphpad Prism v9.3.1. Sample size is n=3 for all numerical data. *srCAR kinetic parameters on TPA are reported as apparent values due to difficulty decoupling monofunctional and bifunctional reduction.

With the aid of the Joint Genome Institute, we cloned the 22 CARs chosen from the SSN into a vector containing the *sfp* gene from *Bacillus subtilis* (pZE/His-CAR-RBS2-*sfp*, see **SI Methods**). Concomitant expression of CAR with *sfp*, which encodes a 4’-phosphopantetheinyl transferase (PPTase), allows for activation of the apo-CAR through the posttranslational addition of a phosphopantetheine arm.^56^ This modification allows for the transfer of the bound thioester intermediate from the adenylation domain to the reduction domain during the catalytic cycle. When expressed under standard conditions, we noticed that several CARs exhibited poor expression. We optimized the media and expression temperature (**Table S1**) and were able to visualize soluble expression of 17 CAR orthologs by performing SDS-PAGE (**Fig. S2**) and a Western blot on the cell lysates (**Fig. 3B**). We were then able to purify all 17 CARs using Ni-affinity chromatography.

After purifying our CAR enzymes, we investigated the *in vitro* activity of our purified CARs on potential PET deconstruction products, starting with TPA (**Fig. 3C**). By adding 5 mM of carboxylate stock in pure DMSO to a buffer containing a final concentration of 10 mM MgCl_2_, 0.5 mM NADPH, 1 mM ATP, 100 mM HEPES pH 7.5, and 1.5 μM of CAR (∼0.2 mg/mL), we measured the initial rates of each CAR at 25 °C spectrophotometrically at 340 nm by tracking the oxidation of the reductive co-factor NADPH.^57^ We found that our highest performing CAR on TPA was srCAR (from *Segniliparus rotundus*), which had an initial rate of 4.80 ± 0.36 mM NADPH_ox_·h^-^1·μM_CAR_^-1^ (**Fig. 3D, Fig. S3**). By normalizing the initial rates of all the CAR orthologs using the activity of srCAR on TPA, we saw that trCAR (from *Trichoderma reesei*), msCAR (from *Mycolicibacterium smegmatis*) and mavCAR (from *Mycobacterium avium*) show 64.5%, 37.5% and 36.2% normalized relative activity, respectively. During the preparation of this manuscript, the first example of a CAR that exhibits activity on TPA was reported.^58^ This report demonstrated activity of mmCAR (from *Mycobacterium marinum*) on TPA in resting *E. coli* whole cells, with no other CARs tested. Intrigued by that result, we performed our own analysis on mmCAR, albeit in an *in vitro* setting. We saw that mmCAR only exhibited 19.8% normalized activity on TPA relative to srCAR (**Fig. 3D**). Curiously, mmCAR had very low activity *in vitro* on 4FBA, even at increasing concentrations of mmCAR (up to 4.5 μM), indicating that 4FBA may be a poor substrate (**Fig. S4**). Analysis of CAR activity allowed us to characterize several “generalist” CARs, like srCAR, mavCAR and msCAR, which had activity on both TPA and 4FBA. We also noticed unique “specialist” CARs in our heatmap, namely from the fungal kingdom. Two of the top three CARs based on activity on 4FBA were ncCAR (from *Neurospora crassa*) and afCAR (from *Aspergillus fumigatus*), and yet they had minimal level activity on TPA. We were also pleasantly surprised by the breadth of reduction performed by trCAR, a previously untested putative aryl acid reductase, which showed soluble expression and reduction activity on the full substrate scope.^59^

We sought to further aid our cascade design by investigating whether CARs could accept an even broader range of starting molecules that could be derived from PET (**Fig. 3C**). Based on the literature, we hypothesized that CARs may prefer aryl substrates that contain ring substituents with greater electron withdrawing propensity.^57^ Therefore, one compound of interest to us was monomethyl terephthalate (mmTPA). Encouragingly, here we succeeded in converting TPA to monomethyl terephthalate (mmTPA) at a yield of 76% using chemical synthesis (**SI Methods**). Additionally, we were able to convert dimethyl terephthalate, which is a product of methanolysis of PET, selectively to mmTPA at 89% yield (**SI Methods**).^60^ A recent paper also shows that mmTPA can be produced from TPA with high selectivity using a carboxyl methyltransferase from *Aspergillus fumigatus*.^61^ When we tested CAR activity on mmTPA, the general trend across the enzyme family was that CARs exhibited high activity on mmTPA. While srCAR only exhibited a 1.7-fold increase in initial activity on mmTPA when compared to its activity on TPA, niCAR (from *Nocardia iowensis*) exhibited a noteworthy 9.9-fold increase in activity compared to its activity on TPA.

Another potential monoester/monoacid product of enzymatic deconstruction of PET is *mono*- (2-hydroxyethyl)-terephthalic acid (MHET). In fact, MHET is in some ways a more accessible enzymatic deconstruction product of PET given that a MHETase is often the second step in published PETase/MHETase cascades that obtain TPA via MHET.^62^ MHET is a particularly interesting candidate substrate given its large size relative to known substrates of CARs. Excitingly, niCAR, mmCAR, srCAR, and trCAR showed between 1.1 to 4.6-fold higher activity on MHET as compared to their respective activities on TPA. Overall, the results observed for the monoesters mmTPA and MHET were encouraging not just for the faster kinetics observed but also because they offered the potential to produce mono-amine products with the esters serving akin to protecting groups.

When screening CARs against potential PET deconstruction products, we were also curious about activity on potential intermediates from our envisioned enzyme cascade for diamine production. The conversion of TPA to pXYL by a CAR and ω-TA cascade would theoretically result in a reaction fork, where 4FBA could either be reduced by CAR to TPAL, or aminated by ω-TA to pAMBA, an amine-containing aromatic carboxylate (**Fig. 1B)**. When we provided pAMBA as a candidate substrate, srCAR and trCAR showed 23.1% and 13.2% normalized relative activity, respectively, with mmCAR showing detectable, but very poor activity of 5.6% normalized relative activity. The low activity of CARs on pAMBA presented potential drawbacks and benefits – it could be a bottleneck on route to pXYL, but it could increase the likelihood of selectively producing pAMBA when that is the desired product.

To better understand the kinetics of our highest performing CAR, we sought to characterize the k_cat_ and K_m_ of srCAR on TPA, 4FBA, mmTPA, and MHET as shown by the Michaelis-Menten plots (**Fig. 3E**). Using the previously described plate reader-based NADPH oxidation assay, we observed the average k_cat,apparent_ and K_m,apparent_ on TPA is 46.9 min^-1^ (95% C.I. 42.5-52.7 min^-1^) and 2.1 mM (95% C.I. 1.7-2.7 mM), respectively (**Fig. 3F**). We report these values as “apparent” as srCAR is a promiscuous enzyme and catalyzes not only the single reduction of TPA to 4FBA, but also the subsequent reduction of 4FBA to TPAL, which makes it challenging to quantify biochemical parameters for only the first reaction. Fortunately, we can analyze just the second reaction in isolation by supplying 4FBA. We observed that the average k_cat_ and K_m_ of srCAR on 4FBA is 30.3 min^-1^ (95% C.I. 28.8 to 31.8 min^-1^) and 0.33 mM (95% C.I. 0.27-0.41 mM), respectively. We also determined the srCAR kinetic parameters for MHET and mmTPA, which interestingly, exhibit approximately one or two orders of magnitude lower K_m_, respectively, than TPA, allowing for higher catalytic efficiency on these substrates (**Fig. 3F**).

### Sequence and Structural Insights of Natural CARs through Homology Modeling and Molecular Docking

To explore whether the predicted protein structures would have patterns that correlate with the experimental observations, we explored computational analyses. We first generated predictions of the 3D structure of the CAR adenylation domains (A-domains) using AlphaFold 2.1.^63^ Predicted structures are deposited as PDB files in Github (https://github.com/KunjapurLab/CAR-Predicted-Structures.git). For niCAR, the previously generated crystal structure was used instead of a prediction (PDB code: 5MSC).^44^ AlphaFold did not generate a high confidence structure for the putative CAR from *Cafeteria roenbergensis*; thus, we omitted this protein from further analysis. We used the predicted structures to compare binding pocket residues, binding pocket size, and substrate interactions via molecular docking.^64^ We identified binding pocket residues based on previous substrate-bound crystal structures and highly conserved signature sequences in the catalytic domains by multiple sequence alignment using Clustal Omega and Jalview.^47,65^ From this analysis, we found that the binding pocket residues are either highly conserved or vary with no clear pattern unique to our best experimentally observed enzymes (**Fig. S5**). However, when we analyzed the average bottleneck radius using the CAVER webtool, we see that several of the CARs that are highly active on our substrate panel have larger predicted adenylation domain binding pocket radii (**Fig. S6A**).^66^

We then explored molecular docking on PET deconstruction products using the AlphaFold predicted protein structures and Autodock Vina. Docking analysis predicted that TPA could bind the predicted enzyme binding pocket of srCAR (**Fig. S6B**). However, molecular docking did not show any observable trend matching that of our experimentally obtained result. Prior studies have suggested the ε-nitrogen atom of a conserved lysine binding pocket residue can form hydrogen bonds with the acid acyl-AMP complex, particularly with the ribose-ring oxygen atom, and that the distance (d1) between the nitrogen atom and oxygen atom of the Lys residue and ribose ring, respectfully, could be a nimble indicator of CAR activity (**Fig. S6C**).^67,68^ However, we did not see a correlation of our experimental results to the obtained d1 values with our docked AMP-acyl TPA. Interestingly, we observed binding of MHET in the predicted srCAR structure, indicating that the bulky ethylene glycol sidechain could be accommodated at position A315 in the binding pocket (**Fig. S7C**). Overall, our computational analysis indicates that much remains to be understood about CAR specificity and that, for the time being, empirical studies such as ours that sample a wide range of CARs and potential substrates are essential towards elucidating the specificity of this important enzyme family.

### Characterization of CAR Activity on PET Deconstruction Products with a Co-factor Regeneration Cascade

Given the activity of CARs on several substrates of interest, we sought to provide higher substrate loading and monitor reaction progress over time. CARs are dependent on ATP and NADPH to catalyze the reduction of acids; however, supplementation of exogenous co-factors becomes more cost-prohibitive at higher substrate loading. For enzymatic ATP regeneration, we expressed a recently discovered type 2-III polyphosphate kinase (PPK2-III) from an unclassified *Erysipelotrichaceae* bacterium (PPK12) that was coupled to CAR to produced aldehydes.^45^ PPK12 converts AMP to ATP using sodium hexametaphosphate as a polyphosphate (polyP) source. Though initially we noticed our reaction stalling using this enzyme, we later discovered that polyP was inhibitory to either our CAR or PPK12 and that optimizing the polyP concentration allowed the CAR reaction to proceed (**Fig. S8**). For NADPH regeneration, we expressed and tested the activity of an engineered glucose dehydrogenase from *Bacillus megaterium* (bmGDH) (**Fig. S9**).^69^ Lastly, the addition of purified inorganic pyrophosphatase from *E. coli* (ecPPase) eliminated accumulation of inorganic pyrophosphate (PP_i_), which is inhibitory to CAR.^70^

After verifying activity of the key regeneration enzymes, we sequentially characterized the effect of bmGDH and PPK12 with srCAR. At our chosen endpoint of 24 h, we observed that the srCAR-catalyzed reduction of 5 mM TPA does not reach completion to TPAL, even with exogenous supplementation of both NADPH and ATP in excess as measured by HPLC-UV (**Fig. 4A, Fig. S10-S11**). However, we noticed that addition of bmGDH resulted in the formation of an over-reduced product, 4-(hydroxymethyl)benzaldehyde (HMB) at 1 mM concentration of NADP^+^. The formation of alcohols is not known to be catalyzed by CAR,^44^ but has been reported in systems in which protein preparations may contain impurities.^71^ Recently, the ability of some CARs to also catalyze overreduction of aldehydes to alcohols was shown.^72^

**Figure 4.**
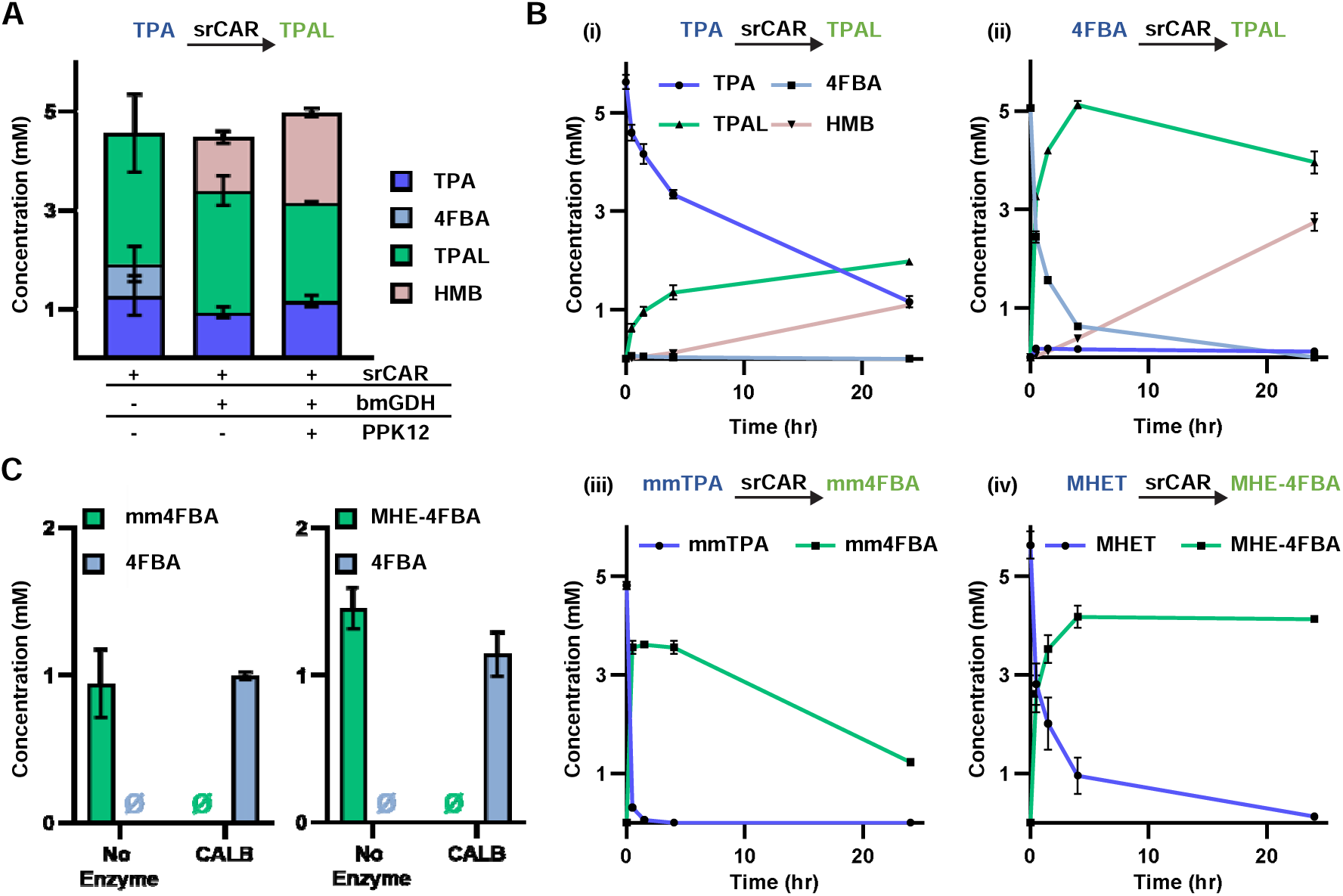
Reduction of PET-derived deconstruction products catalyzed by srCAR. (A) Endpoint *in vitro* assays showing the conversion of 5 mM TPA to TPAL after 24 h, with and without co-factor regenerating enzymes. Our multienzyme cascade shows the overreduction of TPA to an alcohol, 4-(hydroxymethyl)benzaldehyde (HMB). (B) Time course analysis of the enzymatic reduction of (i) 5 mM TPA, (ii) 5 mM 4FBA, (iii) 5 mM mmTPA and (iv) 5 mM MHET by CAR. All experiments included a 3-enzyme NADPH and ATP regeneration system. Reduction of TPA and 4FBA show over-reduction of aldehydes to alcohols. While mmTPA shows a decline in concentration over time to the purported alcohol product, we were not able to independently verify this. To our surprise, MHET shows no over-reduction to the alcohol. (C) At our chosen endpoint of 24 h, lipase from *Candida antarctica* (CALB) was able to completely de-esterified 1 mM of both MHE-4FBA and mm4FBA at 30 °C, providing an alternate route to forming aldehydes from PET-derived esters. Sample size is n=3 and all data are shown as the mean ± standard deviation. Error bars for certain points of the bioconversion of mmTPA to mm4FBA are smaller than the height of the symbol.

To better understand the timescale of CAR catalyzed reduction of PET-derived substrates, we performed a time-course analysis of srCAR-catalyzed reduction of 5 mM TPA, 4FBA, mmTPA, and MHET (**Fig. 4B**). After 24 h, the conversion of TPA is 77%; however, the yield of the desired dialdehyde is only 39%, with 22% of the aldehyde products over-reduced to HMB. Interestingly, by using 4FBA as the starting substrate, we see complete conversion of 5 mM 4FBA to 5 mM TPAL. The improved conversion is consistent with our heatmap analysis and our *in vitro* biochemical kinetic characterization of srCAR, which suggest that the lower substrate K_m_ of 4FBA increases the catalytic efficiency of the reaction. At later timepoints, we again see the undesired production of HMB. Reduction of mmTPA to methyl 4-formylbenzoate (mm4FBA) by srCAR rapidly reached 65% conversion in 30 min but was likely subject to over-reduction, as reported previously.^73^ Fortunately, we observed a potential alternate route to producing aldehydes from PET deconstruction products that mitigated over-reduction. By supplementing 5 mM of MHET to the multienzyme reduction cascade, we saw complete conversion of MHET within 4 h, with a yield of 85% of desired product, 2-hydroxyethyl 4-formylbenzoate (MHE-4FBA).

To verify the viability of this potential alternative reductive pathway to produce aldehydes of interest, we sought to hydrolyze the ester group on both mm4FBA and MHE-4FBA, to form 4FBA, which we showed earlier that srCAR can convert 4FBA to TPAL (**Fig. 4B**). Lipases are well-suited for this application as they perform the hydrolysis of ester bonds in aqueous environments.^74^ Previously, lipase B from *Candida antarctica* (CALB) was used to deconstruct MHET to TPA.^75^ Driven by the structural similarity of these carboxylates to their corresponding aldehydes, we used 1 U of CALB to perform the hydrolysis of 1 mM of both mm4FBA and MHE-4FBA, which yielded 4FBA as a sole product as quantified by HPLC-UV after a 24 h incubation at 30 °C (**Fig. 4C**). The ease of accessing diverse monoester derivatives of terephthalate from PET deconstruction coupled with the broad reduction activity of srCAR, allows for the design of modular enzyme cascades to synthesize two distinct aldehydes, 4FBA or TPAL, and opportunities for downstream functionalization by cvTA.

### One-pot Synthesis of Diamine by a Reductive Biocatalytic Cascade

After determining the optimal CAR and ω-TA for use in our cascade, we developed a one-pot assay to produce our diamine of interest, pXYL. We combined srCAR, cvTA, and the 3-enzyme co-factor regeneration system in a one pot assay with 5 mM TPA (**Fig. 5A, SI Methods**). Using HPLC-UV (**Fig. S12, SI Methods**), we were able to discern all the different potential reaction intermediates of our system. At 1.5 µM loading of both the srCAR and cvTA, we were able to obtain a yield of 35% pXYL and 28% pAMBA, with no aldehyde species present during the reaction endpoint. Encouraged by this result, we looked at tuning the enzyme loading to see if we could shift the product distribution to prefer the diamine. Increasing cvTA concentration from 1.5 µM to 3 µM in the presence of 1.5 µM of srCAR slightly shifted product yield to 38% pAMBA and 26% pXYL. Upon addition of 3 µM of srCAR, we saw a shift in distribution to favor the diamine. Our highest yield in this experiment was 62% yield of pXYL at a 3 µM loading of srCAR and 1.5 µM loading of cvTA. Coupling srCAR to cvTA resulted in no detectable formation of alcohols, which we observed when srCAR was coupled only to the cofactor regeneration cascade

**Figure 5.**
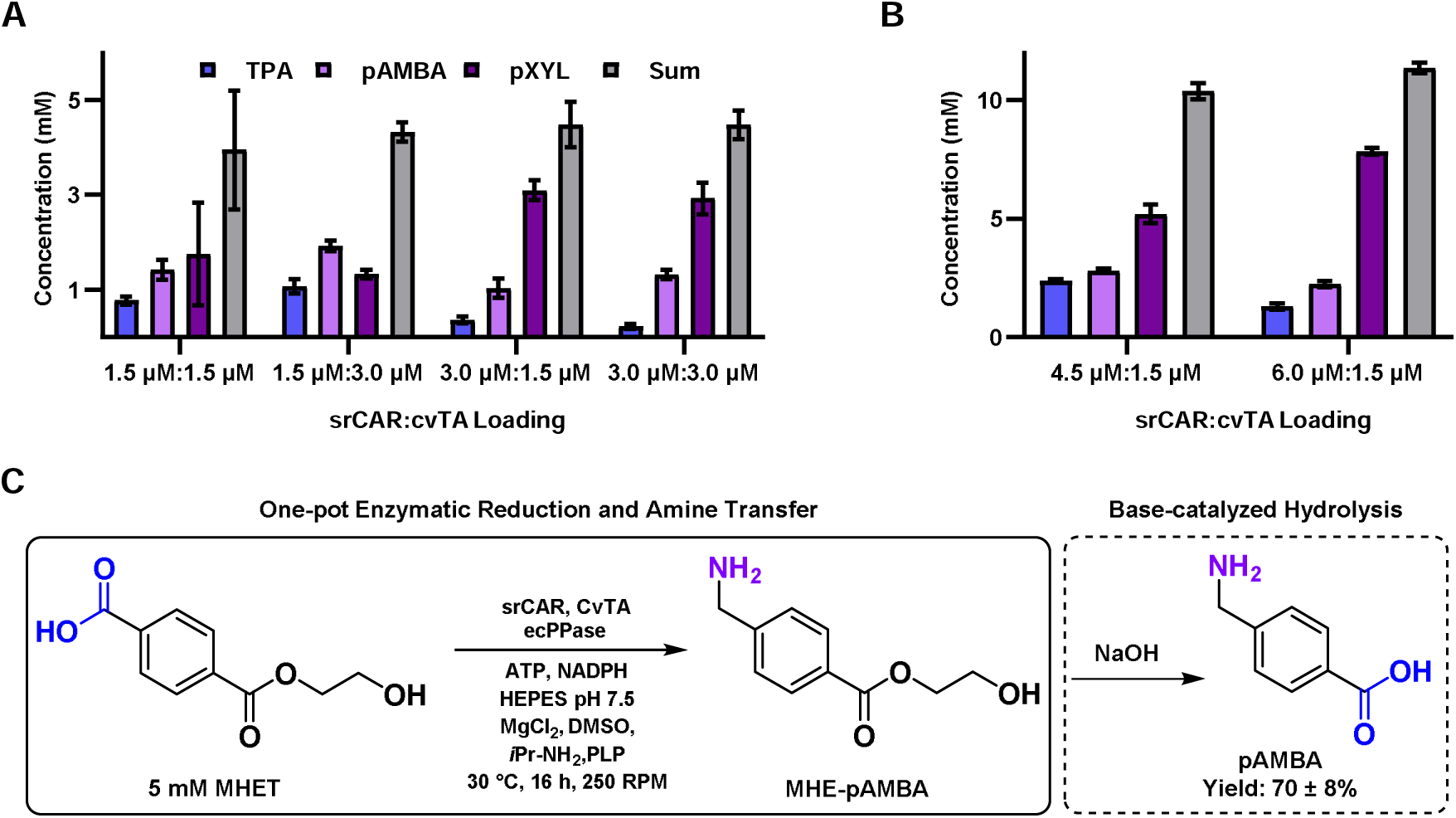
One-pot bioconversion of PET-derived carboxylates to amines. (A) One-pot *in vitro* bioconversion of TPA to pXYL by a 5-enzyme cascade consisting of srCAR, cvTA and a 3-enzyme co-factor regeneration cascade yields high reaction selectivity for amines and an appreciable yield of 62% of the desired diamine pXYL at 3 µM srCAR loading. By tuning the ratio of CAR to TA, we see a corresponding increase in yield and selectivity towards the diamine when CAR concentration is increased. (B) At 10 mM loading of TPA as substrate, we demonstrate 69 ± 1% yield of TPA to pXYL in one-pot at a srCAR:cvTA ratio of 6.0 µM:1.5 µM. (C) The substrate specificity of srCAR and cvTA allows us to produce the pharmaceutically relevant monoamine, pAMBA, solely by switching the starting substrate from TPA to MHET, and performing a simple hydrolysis step to liberate pAMBA, yielding 70 ± 8% of the desired product. For the bioconversion of MHET to pAMBA, 6.0 µM srCAR, 1.5 µM cvTA (dimer basis), 0.75 µM ecPPase (hexamer basis), 6 mM NADPH, 6 mM ATP, 20 mM MgCl_2_, 100 mM HEPES pH 7.5, 5% (v/v) DMSO, 20 mM *i*Pr-NH_2_, 400 µM PLP, and 5 mM of MHET were added to a total volume of 100 µL. Following a 16 h incubation at 30 °C, the reaction was basified by NaOH to a final concentration of 0.5 M and analyzed using HPLC-UV. Sample size is n=3 and all data are shown as the mean ± standard deviation.

To test the cascade performance at higher substrate loading, we increased loading of TPA to 10 mM. From our previous experiments, we saw that the increase in srCAR concentration drives the product distribution towards pXYL. We therefore further increased the concentration of srCAR to 4.5 µM and 6.0 µM, keeping the concentration of cvTA at 1.5 µM. We achieved a 69 ± 1% yield of our target diamine at 6.0 µM srCAR and 1.5 µM cvTA loading (**Fig. 5B**). This result is unusually promising given the presence of only one observed by-product, pAMBA, at 22% of the final reaction mix, the relatively low concentration of enzyme, and the ability to use single variants of CAR and TA enzymes. Our enzyme cascade achieves a higher yield of desired diamine (69% instead of 30%) than a previously reported CAR/TA cascade that was supplied with 10 mM adipic acid.^40^ There are several differences between these two efforts that are worth noting. Our system features aromatic compounds as opposed to medium-chain aliphatic compounds, thus generating aldehyde intermediates that differ with respect to reactivity and volatility. Our system for diamine production features the provision of lower cost precursors to required CAR co-factors (AMP and NADP^+^) instead of the actual CAR co-factors (ATP and NADPH). Our system also includes a different amine donor (we use *i*Pr-NH_2_ instead of alanine) and no regeneration of the amine donor. Finally, our cascade features reduced complexity by identifying single CAR and TA variants capable of performing the bifunctional reduction and amination reactions. However, despite these differences in conditions, our enzyme concentrations were similar (6 μM srCAR in our cascade vs. 3 μM each of two CAR variants).

### One-pot Synthesis of Monoamine by a Reductive Biocatalytic Cascade

Lastly, we took advantage of the wide substrate scope of both srCAR and cvTA to design an enzyme cascade for the synthesis of monofunctionalized amine, pAMBA, an antifibrinolytic drug.^26^ We had previously observed selective reduction of MHET to MHE-4FBA by srCAR (**Fig. 3D, Fig. 4B)**. Owing to the wide substrate scope of cvTA, and the previous result that cvTA is highly active on a structurally similar aldehyde, 4FBA, we posited that MHE-4FBA can also be accepted as a substrate by cvTA. In a one-pot reaction with 6.0 µM srCAR, 1.5 µM cvTA, and 5 mM of MHET as the starting substrate, we observed the formation of an amine peak via HPLC-UV after performing an amine-specific derivatization (**SI Methods**). Encouraged by this result, we then performed base-catalyzed hydrolysis, yielding pAMBA at 70 ± 8% yield, with no diamine formation (**Fig. 5C).** The wide substrate scope of the CAR and ω-TA allows for a modular route to upcycled two distinct amine products, with high selectivity, depending on the available PET-derived starting substrate.

## Conclusion

New technologies must be developed to combat the growing amount of plastic waste, and chemical deconstruction methods alone may not sufficiently incentivize nor diversify the recycling of plastic waste. Biocatalytic synthesis of diamines and monoamines from PET-derived esters or carboxylic acids monomers offers an opportunity upcycle PET for applications in materials and therapeutics. Given the selectivity of enzymes, one would expect that the enzymes in our cascade would be amenable to relatively seamless integration with enzymatic deconstruction approaches, such as those that rely on cutinases or PETases. In principle, this integration could also occur in one pot, increasing the attractiveness and cost-competitiveness of biocatalytic valorization strategies. Additionally, because the aromatic character of the starting material is preserved and only a small number of molecular transformations are required, the routes to these products offer higher theoretical yields than when starting from traditional fermentative carbon sources such as glucose.

This study revealed several interesting aspects of substrate specificity for three industrially relevant enzyme classes. We first established that cvTA could catalyze amine transfer to the PET-derived aldehydes 4FBA and TPAL, which resulted in a high yield of the desired amines pXYL and pAMBA, respectively. With this knowledge, we then explored enzymatic synthesis of aldehydes from PET-derived substrates. Through a bioprospecting approach, we identified that srCAR could catalyze the bifunctional enzymatic reduction of TPA to TPAL (via 4FBA), which is a relatively newly observed phenomenon. Moreover, it was encouraging that this single CAR could accept MHET and mmTPA as substrates for reduction to their respective monoaldehyde products, which we show could be de-esterified enzymatically by a lipase. Our bioprospecting approach also revealed potential utility of a previously uncharacterized putative aryl acid reductase from *Trichoderma reesei*, whose activity and substrate scope were favorable and merit further testing.

By pairing enzymatic aldehyde production to amine transfer, we developed a one-pot route to both produce diamines and mono-amines. Upon addition of 10 mM TPA to srCAR, cvTA and enzymes for co-factor regeneration, we achieved 69 ± 1% of the desired diamine pXYL. We also highlight the ability of our chosen enzymes to accept multiple starting substrates depending on the desired product output. This modularity is highlighted in the one-pot enzymatic reduction and amine transfer of MHET, which, followed by base-catalyzed ester hydrolysis, produced the monoamine therapeutic pAMBA at high selectivity and yield of 70 ± 8%, from 5 mM of the starting substrate. Given that only micromolar enzyme concentrations were used, we believe this cascade has potential for polymer and medicinal synthesis, where larger quantities and purities of amine products would be required. Overall, our reductive cascade design aids in further incentivizing the diversion of PET waste from the environment by augmenting the sparse list of biological PET upcycling strategies.

## ASSOCIATED CONTENT

### Supporting Information

The following files are available free of charge. Supplementary Methods, Figures (S1-S12) and Tables (S1-S3) (PDF)

## AUTHOR INFORMATION

### Author Contributions

The manuscript was written through contributions of all authors. All authors have given approval to the final version of the manuscript.

### Notes

A.M.K., M.R.G., R.M.D., and W.C. have filed for a provisional patent related to this work.

## ACKNOWLEDGMENT

The authors would like to thank the members of the University of Delaware Center for Plastics Innovation for guidance and support on this project.

## ABBREVIATIONS

CAR: carboxylic acid reductase
ω-TA: ω-transaminase
PET: polyethylene terephthalate
TPA: terephthalic acid
4FBA: 4-formylbenzoic acid
TPAL: terephthalaldehyde
pAMBA: *para*-(aminomethyl)benzoic acid
pAMB: *para*-(aminomethyl)benzaldehyde
pXYL: *para*-xylylenediamine
HMB: 4-(hydroxymethyl)benzaldehyde
MHE-4FBA: 2-hydroxyethyl 4- formylbenzoate
mm4FBA: methyl 4-formylbenzoate
BHET: *bis*-(2-hydroxyethyl) terephthalate
MHET: *mono*-(2-hydroxyethyl) terephthalic acid
NADPH: nicotinamide adenine dinucleotide phosphate (reduced)
NADP^+^: nicotinamide adenine dinucleotide phosphate (oxidized)
ATP: adenosine 5′-triphosphate
AMP: adenosine 5′-monophosphate

