## Supplemental Information for "Reductive Enzyme Cascades for Valorization of PET Deconstruction Products"

Supporting Information

Madan R. Gopal[1,3], Roman M. Dickey[1,3], Neil D. Butler[1], Michael R. Talley[2,3], Ashlesha Mohapatra[1], Mary P. Watson[2,3], Wilfred Chen[1,3], Aditya M. Kunjapur[1,3*]

**Affiliations:**

[1]Department of Chemical & Biomolecular Engineering, University of Delaware, Newark, DE 19716

[2]Department of Chemistry and Biochemistry, University of Delaware, Newark, DE 19716

[3]Center for Plastics Innovation, University of Delaware, Newark, DE 19716

**Supporting Information: Table of Contents**

#### Supplemental Figures

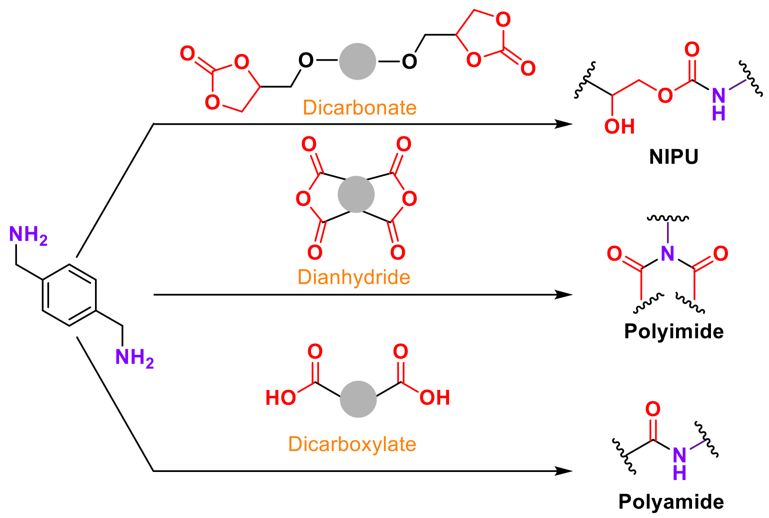

**Figure S1.** Diamines like pXYL are used as monomers in various thermoset and thermoplastic polymers. Shown here are three potential polymer applications for pXYL. pXYL can be reacted with various comonomers can be used to form nonisocyanate polyurethanes (NIPU), polyimides and polyamides.

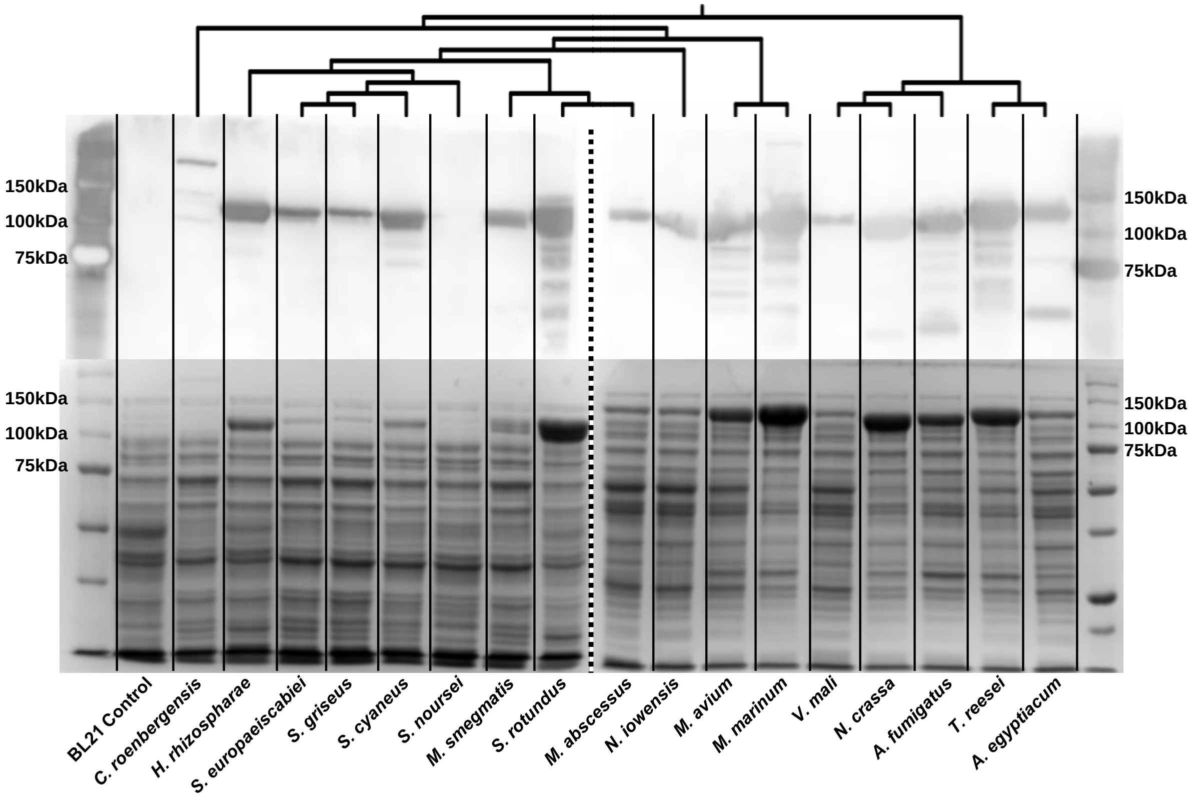

**Figure S2.** Anti-His$_6$ Western (top) and SDS-PAGE (bottom) images of CAR expression ordered by phylogeny as reported by Clustal Omega sequence alignment.[1] The phylogenetic tree is rooted at the midpoint and ignores branch lengths. CAR lysate was denatured in SDS and β-mercaptoethanol prior to loading 11.25 μg total protein onto a 10% SDS-PAGE gel. The SDS-PAGE and Western blot lanes came from the same lysate samples loaded onto identical 10% SDS-PAGE gels. Cells were grown under optimal expression conditions and therefore relative expression cannot be compared by the Western blot.

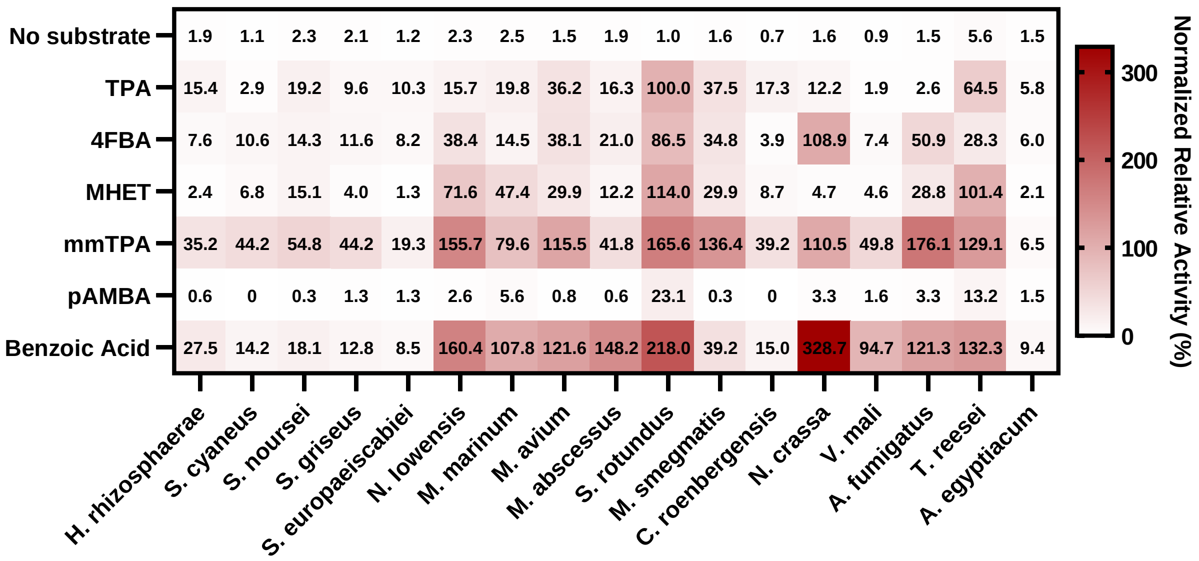

**Figure S3.** Here we report the full activity values of all 17 CAR orthologs on TPA, 4FBA, MHET, mmTPA, pAMBA and the model aromatic acid, benzoic acid. The heatmap values represent the average initial rate, normalized to the average activity of srCAR on TPA of 4.80 mM $NADPH_{ox} \cdot h^{-1} \cdot \mu M_{CAR}^{-1}$. Heatmap cells below 5% normalized relative activity were considered to have no detectable activity. Data is shown as the average of triplicates.

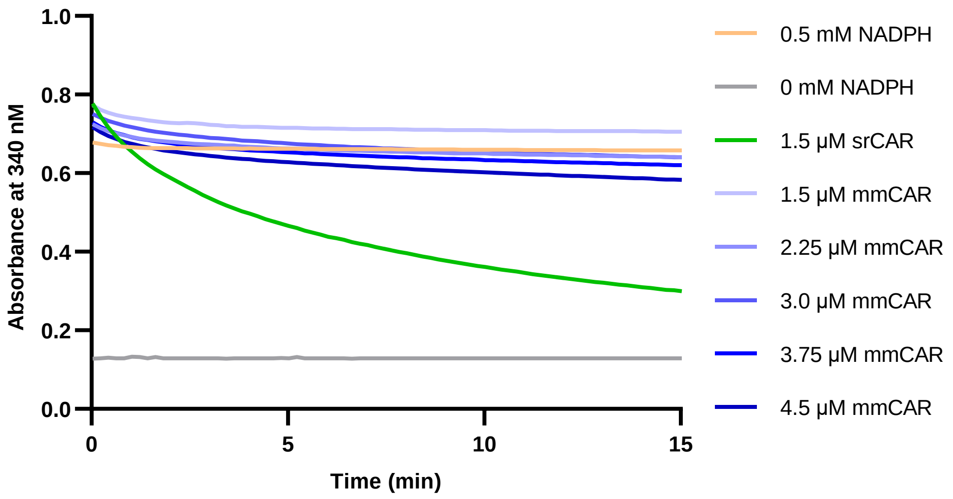

**Figure S4.** *In vitro* CAR assay to evaluate mmCAR and srCAR as catalysts for the conversion of 4FBA to TPAL. After verifying protein concentration by Bradford assay normalized to a BSA standard curve, we loaded 1.5 µM of srCAR, and increasing concentrations of mmCAR (1.5 µM to 4.5 µM) We observed that even at the higher enzyme loading, mmCAR was not able to reduce 5 mM 4FBA compared to 1.5 µM srCAR, as measured by the oxidation/depletion of the reductive cofactor NADPH at 340 nm. Assay was run simultaneously for both enzymes following the in vitro kinetic assay procedure (see ***In vitro* CAR Kinetic Activity Assay**). Data is shown as the average of triplicates.

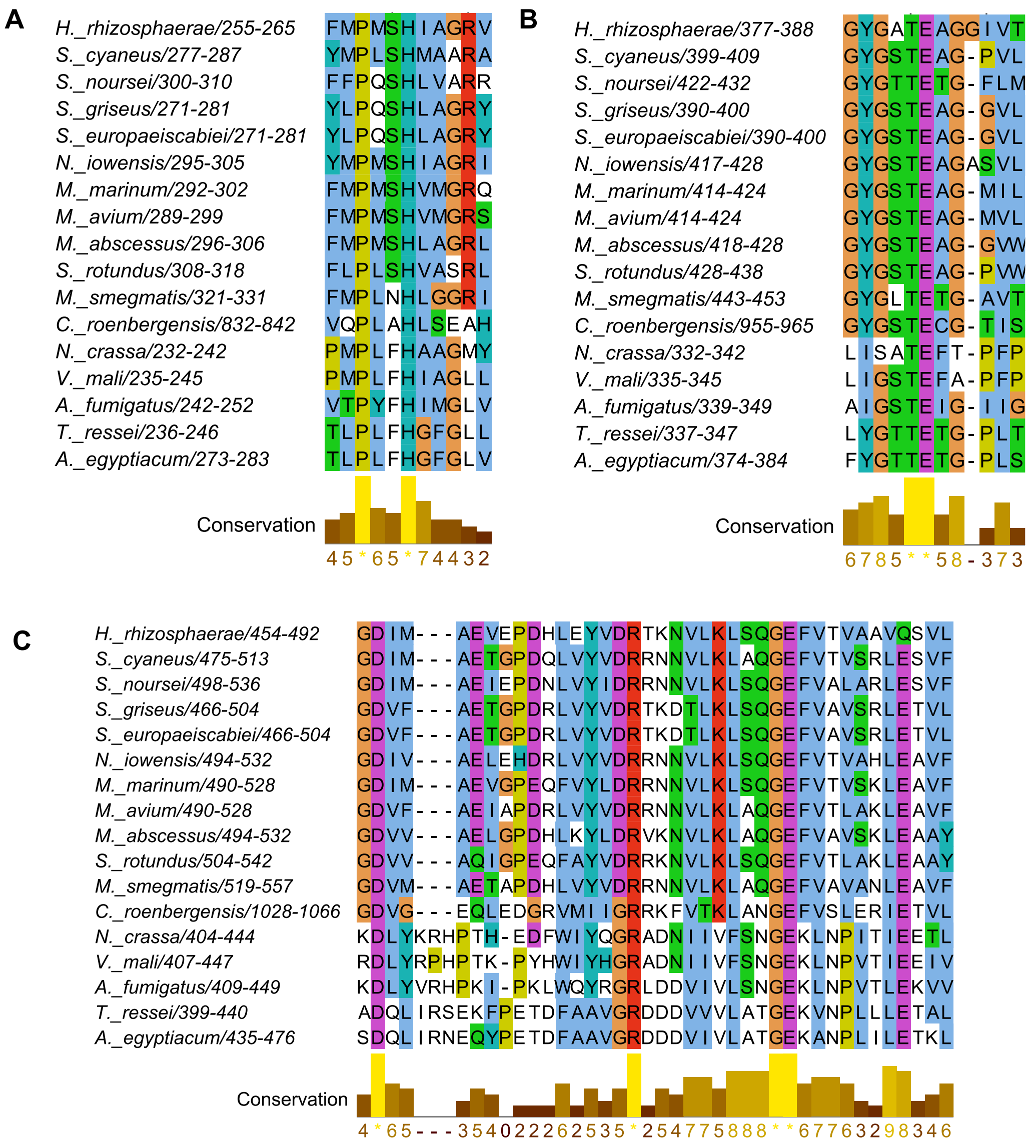

**Figure S5.** Multiple sequence alignment of binding residues identified via previously generated crystal structures and conserved sequences in the catalytic domain of CARs (blue: hydrophobic, red: positively charged, magenta: negatively charged, green: polar, orange: glycine, yellow: prolines and cyan: aromatic). (**A**) Sequence with predicted function of substrate binding. (**B**) Sequence with predicted function of AMP binding. (**C**) Sequence of predicted function of AMP and substrate binding.

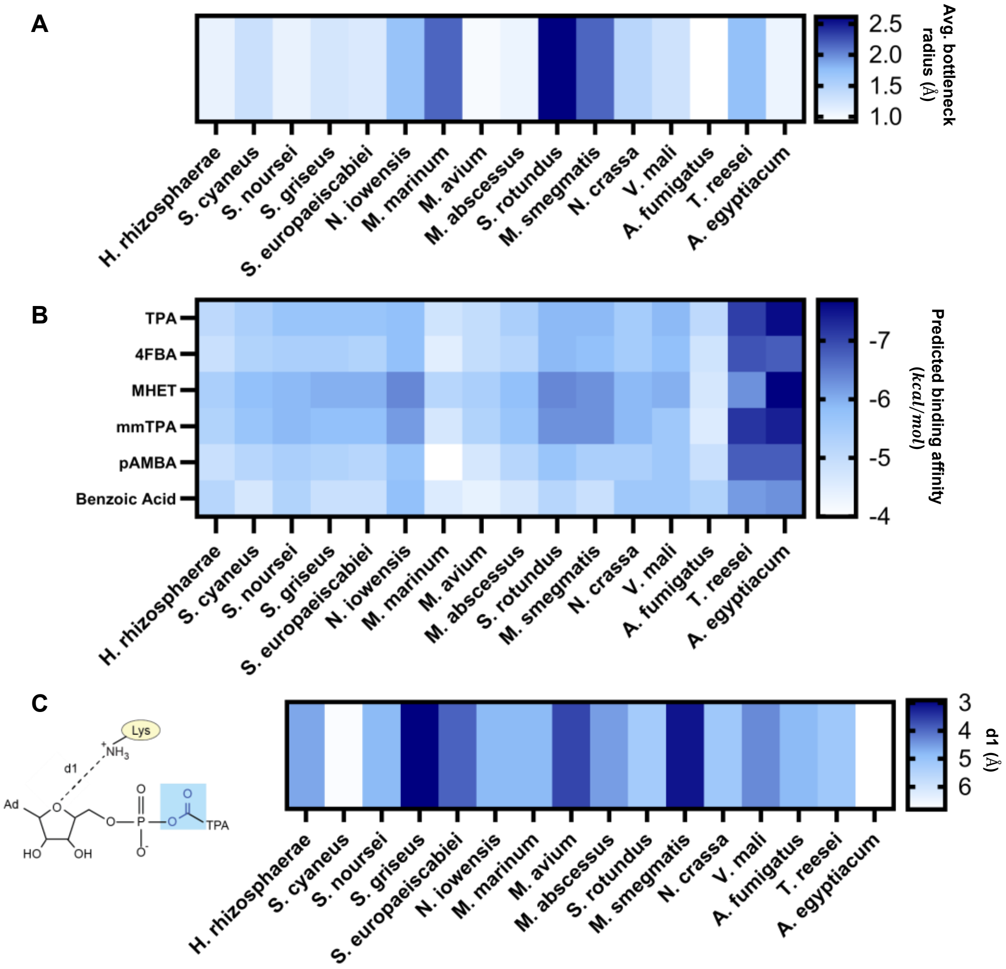

**Figure S6.** Alphafold was used to generate the A domain of CARs without a known crystal structure. Crystal structure PDB: 5MSC from *Nocardia iowensis* was used instead of predicted structures. AlphaFold did not generate a high confident structure for *Cafeteria roenbergensis*. **(A)** Average bottleneck radius of our previously generated CAR structures. Average bottleneck radius was calculated using the CAVER online tool by selecting conserved binding pocket residues of the CARs. **(B)** Heat map of predicted binding affinity of TPA, 4FBA, MHET, mmTPA, pAMBA and benzoic acid. Predicted binding affinity was found using Autodock Vina and Chimera. **(C)** The distance (d1) generated from docking of an acyl-AMP TPA compounds, where d1 is the distance between the ε-nitrogen atom of a conserved lysine binding pocket residue and the ribose-ring oxygen atom in acid acyl-AMP compound.

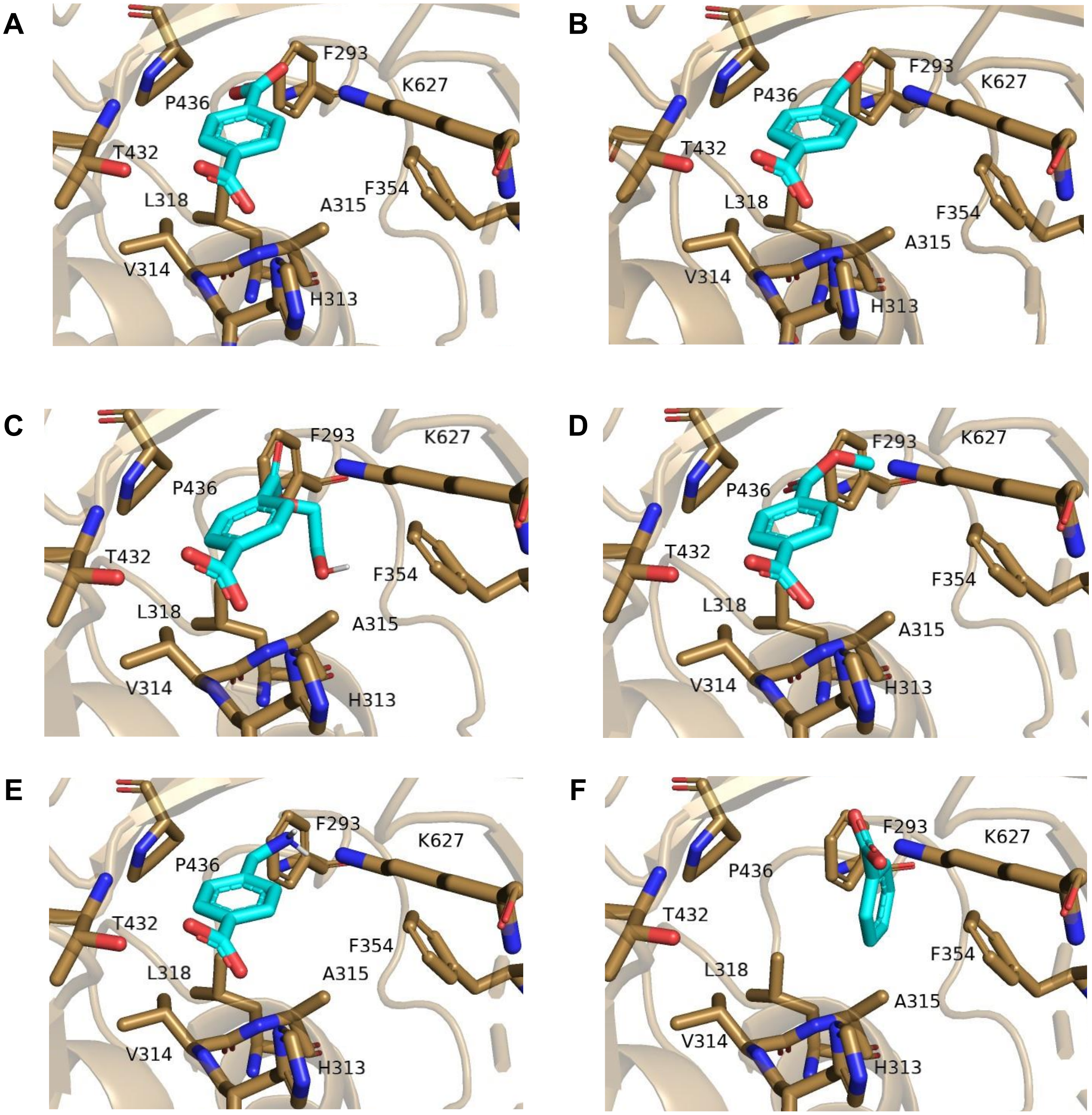

**Figure S7. :** Predicted docking of substrates with srCAR using Autodock Vina **(A)** TPA, **(B)** 4FBA, **(C)** MHET, **(D)** mmTPA, **(E)** pAMBA **(F)** benzoic acid

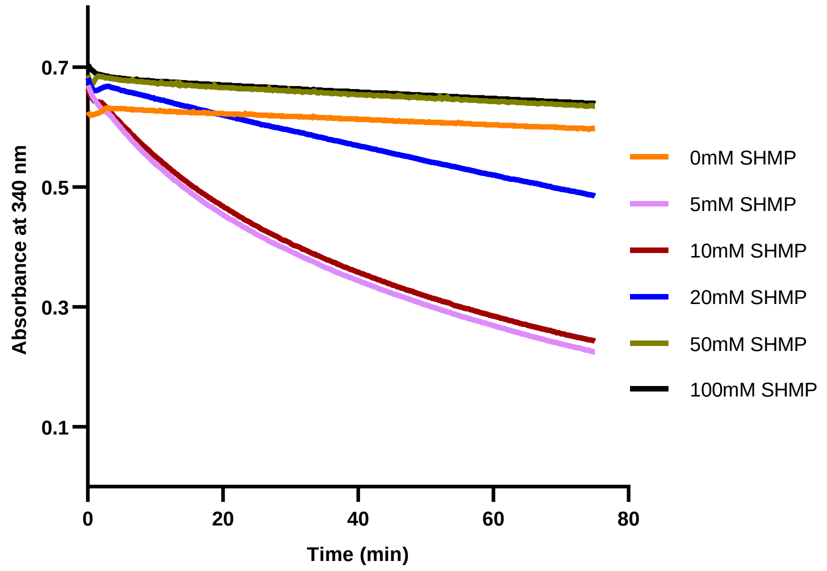

**Figure S8.** After purification of the polyphosphate kinase 2-III (PPK12), an in vitro activity assay was performed to test its activity. 1.5 µM mavCAR and 750 nM ecPPase (hexamer basis) were added to 0.05 mM of ATP, 0.5 mM NADPH (Santa Cruz Biosciences, tetrasodium salt), 10 mM $MgCl_2$, 100 mM HEPES pH 7.5, 5% v/v DMSO, 5 mM benzoic acid, 50 mM glucose (no bmGDH added), and varying concentrations of sodium hexametaphosphate (SHMP, Thermo Scientific). After mixing the reaction components into a 96-well plate, followed by 15 min benchtop incubation to allow the CAR reaction to consume the exogenously supplemented 0.05 mM ATP, 0.42 mg/mL of purified PPK12 was added to the reaction, allowing the AMP produced by the CAR reaction to be regenerated to ATP. The newly regenerated ATP and exogenously supplemented NADPH provides the CAR with the necessary cofactors to perform reduction of benzoate, resulting in a depletion in NADPH which was measured at 25 °C for triplicate samples using an Agilent BioTek Synergy H4 Hybrid Microplate Reader at a wavelength of 340 nm for 75 min. Data is presented as the triplicate average.

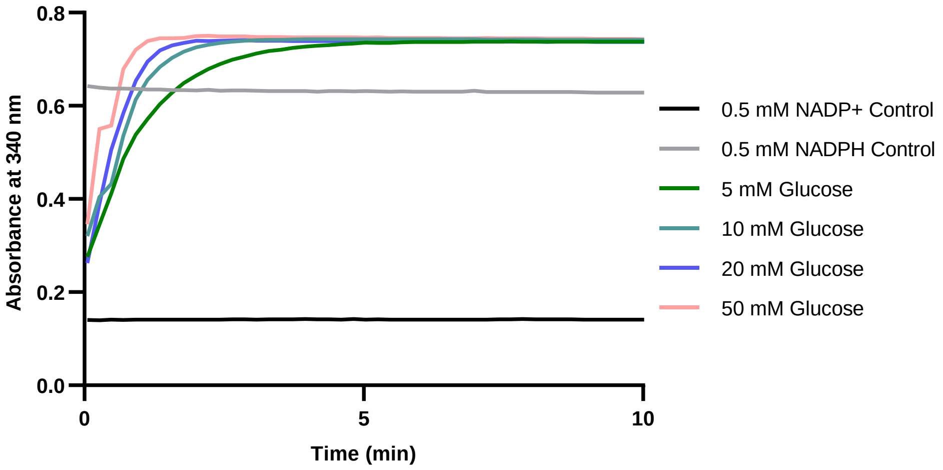

**Figure S9.** After purifying the glucose dehydrogenase (bmGDH), we designed an in vitro assay to test its activity to regenerate NADPH from $NADP^+$ and glucose. 250 nM of bmGDH (tetramer basis), was added to 0.5 mM $NADP^+$ (Santa Cruz Biotechnology, monosodium salt), 100 mM HEPES pH 7.5, and varying concentrations of glucose (5 mM - 50 mM). The reaction was loaded onto a 96-well plate reader at a volume of 100 µL in triplicate. The reduction of $NADP^+$ to NADPH was measured at 25 ºC for triplicate samples using an Agilent BioTek Synergy H4 Hybrid Microplate Reader at a wavelength of 340 nm for 10 min. Reduction of $NADP^+$ to NADPH is shown through an increase in absorbance over time. Data is presented as the triplicate average.

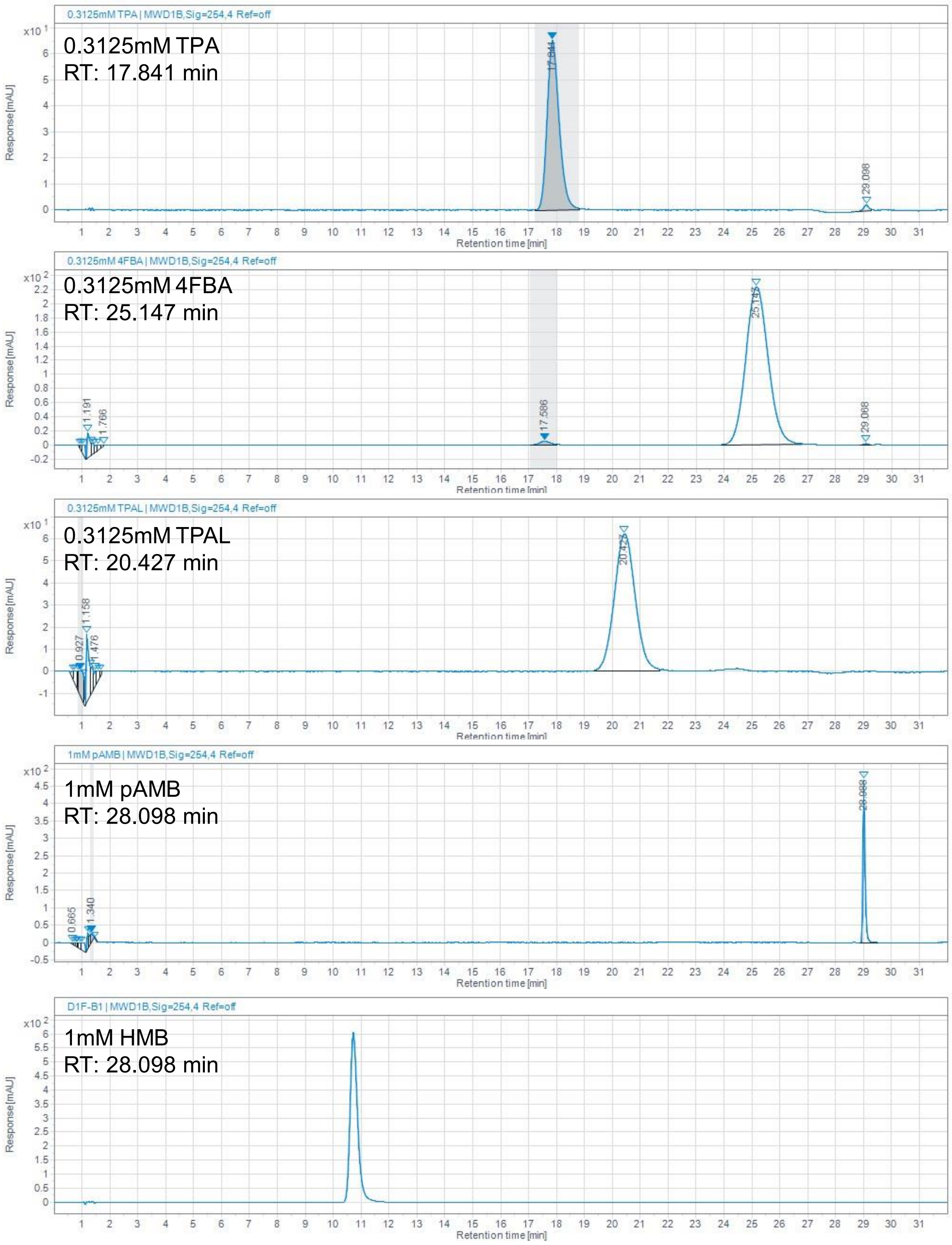

**Figure S10.** Using HPLC Method 1, we show the separation of TPA, 4FBA, TPAL, pAMB, and HMB. The method showed reproducible separation of these compounds. The method is described in detail in Supplemental Methods.

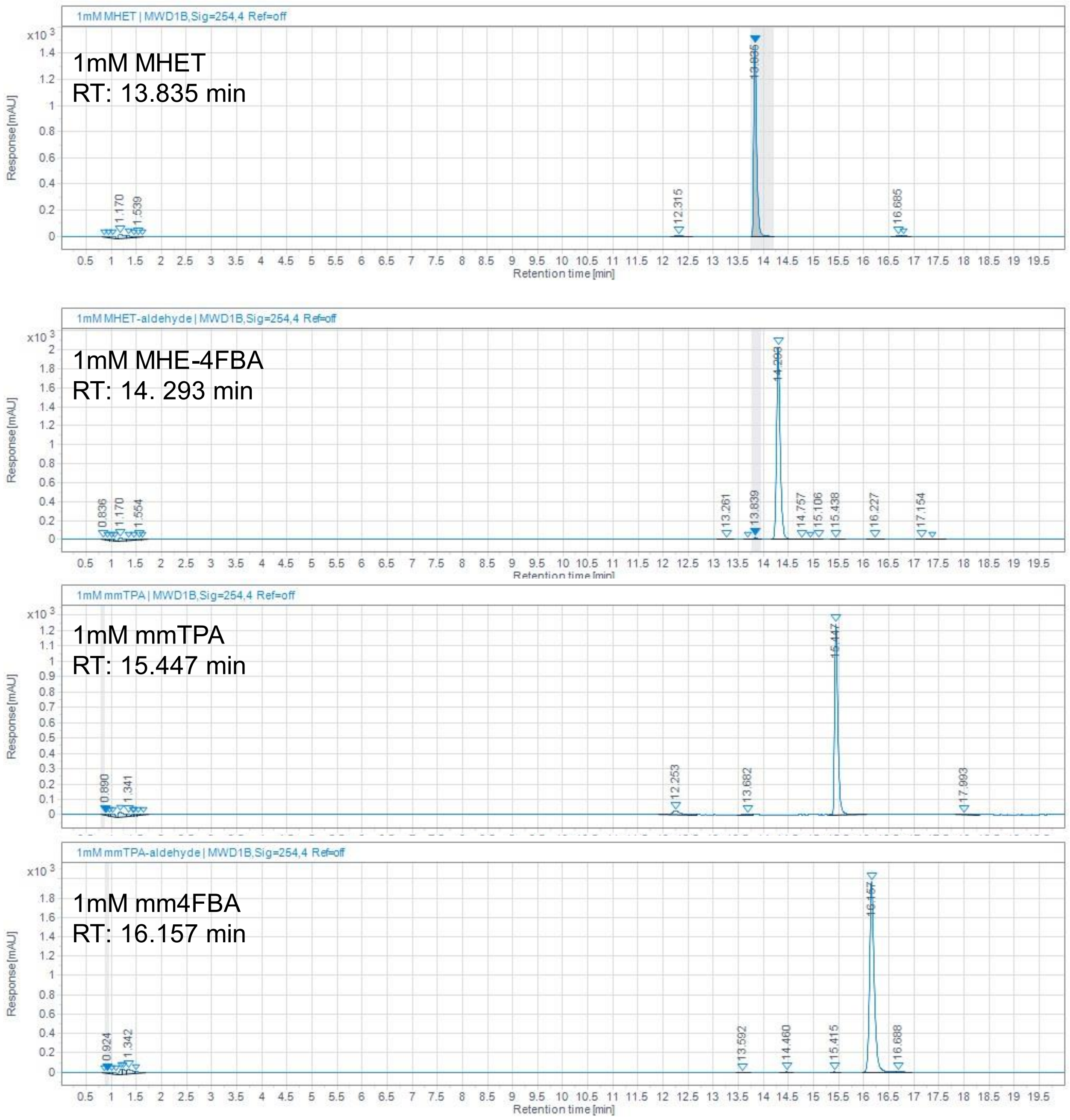

**Figure S11.** Using HPLC Method 2, we show separation of MHET, MHE-4FBA, mmTPA and mm4FBA. The method showed reproducible separation of these compounds. The method is described in detail in Supplemental Methods.

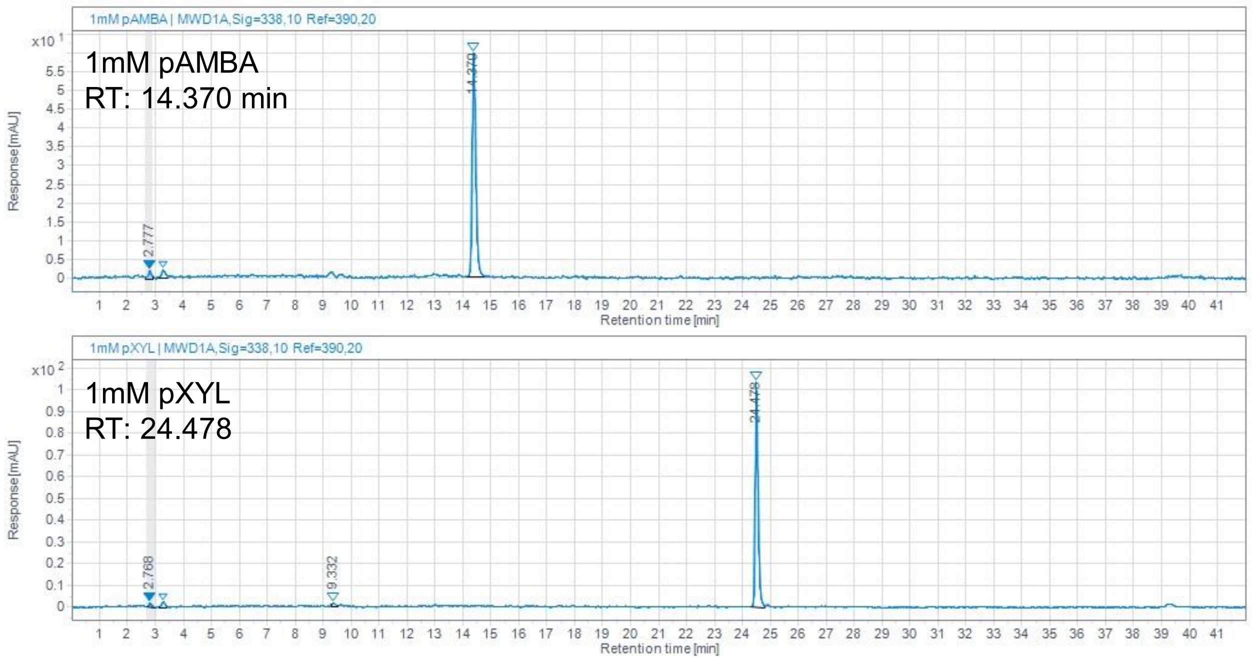

**Figure S12.** Using HPLC Method 3, we show separation pAMBA and pXYL. The method showed reproducible separation of these compounds. The method is described in detail in Supplemental Methods.

### Supplemental Methods

#### Computational Methods

##### Curation of Sequence Similarity Network

To generate the sequence similarity network, we performed NCBI BLAST search to obtain the 500 most closely related sequences to each of these CARs as measured by BLASTP. After deleting duplicate sequences, we obtained 2012 unique sequences, which we then submitted to the Enzyme Function Initiative-Enzyme Similarity Tool (EFI-EST)[2] to generate a sequence similarity network (SSN). To finalize the SSN, we selected an alignment score edge cutoff of 200 and created a 95% representative node structure. Visualization of the SSN using Cytoscape revealed clusters with groupings relatively according to type.

##### Computational Prediction and Sequence Alignment Analysis

The 3D predicted protein structure of the A domain of CARs without a known crystal structure were generated using AlphaFold 2.1. The crystal structure for the CAR A domain from *Nocardia iowensis* (PDB code: 5MSC) was used instead of predicted structures. The resulting protein structures were further used for docking experiments. Substrates, including the acyl-AMP intermediates of TPA, were docked into the active site with 5MSC as a reference using Autodock Vina and Chimera. Average bottleneck radius of predicted structures was calculated using CAVER by selecting highly conserved binding pocket residues of the CARs.

#### Cloning and Purification

##### Cloning, Expression and Purification of CAR

Molecular cloning and vector propagation were performed in *E. coli* DH5α (NEB). CAR genes from *Mycobacterium avium, Trichoderma reesei, Neurospora crassa, Segniliparus rotundus, Mycobacteroides abscessus, and Nocardia iowensis* were purchased as clonal genes from Twist Bioscience and codon optimized expression in for *E. coli*. Codon-optimized CARs were PCR amplified using either NEB Q5 Hot Start Polymerase or KOD Xtreme polymerase. All primers were ordered from IDT. Amplified fragments were then cloned into a PCR-amplified pZE vector harboring an N-terminal $His_6$-tag for purification, kanamycin resistance gene for antibiotic selection, ColE1 ori, and a recombinant 4'-phosphopantetheinyl transferase gene (*sfp*) from *Bacillus subtilis.* Both CAR and *sfp* were placed under the control of the pLtetO for concomitant expression. *Sfp* is required for post-translational activation of the apo-CAR to the holoenzyme. The remaining 16 CARs were cloned by Golden Gate assembly. All plasmids were verified by Sanger sequencing.

For expression, all CAR plasmids were transformed into *E. coli* BL21(DE3) (NEB). 250 mL of LB in 1 L baffled flasks were supplemented with 50 µg/mL kanamycin (Fisher Scientific) was inoculated from a saturated overnight culture to an initial OD600 of 0.1 and induced at mid-exponential phase (OD600 0.5-0.8) with 0.1 µg/mL anhydrotetracycline (ATC, Cayman Chemical). The media type and conditions after induction are listed in **Table S1**. Cells were harvested by centrifugation.

For purification, cells were resuspended in Ni-binding buffer (100 mM HEPES pH 7.4, 250 mM NaCl, 10 mM imidazole, 10% glycerol, 10 mM $MgCl_2$). The cells were lysed by sonication followed by centrifugation at 48,000 × g for 1 h. The cleared supernatant was sterile filtered through a 0.22 µm syringe filter and purified using an AKTA Pure fast protein liquid chromatography (FPLC) system containing a Ni-Sepharose affinity chromatography (HisTrap HP, 5mL). After performing an isocratic wasg at 25 mM and 50 mM imidazole, elution was performed at 250 mM imidazole. Purified fractions were pooled, concentrated, and dialyzed against dialysis buffer (100 mM HEPES pH 7.5, 300 mM sodium chloride, 10% glycerol and 10 mM $MgCl_2$) with a 30 kDa molecular weight cutoff centrifugal filter (Amicon Ultra, Millipore). Samples were then flash frozen in Eppendorf tubes in an ultracold ethanol bath and stored at -80 ºC.

**Cloning, Expression and Purification of *Chromobacterium violaceum* ω-Transaminase, Polyphosphate Kinase 12, *E. coli* Inorganic Pyrophosphatase, and *Bacillus megatarium* Glucose Dehydrogenase**

The ω-transaminase from *Chromobacterium violaceum* (cvTA) and polyphosphate kinase 12 (PPK12) were purchased as gene fragments from IDT and cloned using Gibson Assembly into a pACYC vector harboring an N-terminal $His_6$-tag for purification, p15a ori, and chloramphenicol acetyltransferase gene for antibiotic selection. The inorganic pyrophosphatase gene was amplified by PCR from the *E. coli* NEB5α genome and was cloned into the same pACYC by Gibson Assembly. The glucose dehydrogenase from *Bacillus megatarium* (bmGDH, pET-28a(+)-GDH) was a gift from Frank Schulz (Addgene plasmid # 108954)[3].

All plasmids were transformed into *E. coli* BL21(DE3) (NEB). 250 mL of LB in a 1 L baffled flask was supplemented with appropriate antibiotic. The flask was inoculated from a saturated overnight culture to an initial OD600 of 0.1 and induced at mid-exponential phase (OD 0.5-0.8) with 1 mM isopropyl ß-D-1-thiogalactopyranoside (IPTG, Fisher Scientific). The media type and conditions after induction are listed in **Table S1**. Cells were harvested by centrifugation.

To purify enzymes, cells containing bmGDH, ecPPase, and PPK12 were resuspended in Ni-binding buffer. Cells containing cvTA were resuspended in Ni-binding buffer with 400 μM pyridoxal 5'-phosphate (PLP, TCI America). Cells were lysed by sonication and centrifuged at 48,000 × g for 1 h. The cleared supernatant was sterile filtered through a 0.22 μm syringe filter and was purified using an AKTA Pure fast protein liquid chromatography (FPLC) system and Ni-Sepharose affinity chromatography (HisTrap HP, 5mL) using the same purification protocol as the CAR enzyme (see **Cloning, Expression and Purification of CAR**).

**SDS-PAGE and Western Blot Analysis of CAR Expression**

5 mL $LB_{kan}$ was inoculated with *E. coli* BL21 (DE3) cells harboring CAR plasmids and grown at 37 °C overnight. Cells were seeded at an initial OD600 of 0.1 in 50 mL of media containing kanamycin. An *E. coli* BL21 (DE3) negative control harboring no plasmid was seeded in LB. The media type and conditions after induction are listed in **Table S1**. Cells were then harvested by centrifugation and lysed using silica beads. After clearing cell debris by centrifugation, total lysate protein concentration was measured by a Bradford Assay, using bovine serum albumin (BSA) as calibration standard. Total protein from each CAR lysate was denatured in SDS and β-mercaptoethanol. 2.11.25 μg of total protein was then loaded onto a 10% SDS-PAGE gel and run at 150 volts for 75 min. Gels destined for Western Blot were referenced against a prestained ladder (Bio-rad cat #1610373) while those used for SDS-PAGE analysis of total protein were referenced against an unstained ladder (Bio-rad cat #1610363) and stained with Brilliant Blue.

For Western blot analysis of lysate, proteins from SDS-PAGE were transferred to a Immobilon-E polyvinylidene difluoride membrane (EMD Millipore). The membrane was incubated with 5% fat free milk in Tris-buffered saline with 0.1 μg/mL anti-$His_6$ antibodies (Proteintech). Chemiluminescent reagents (Amersham ECL Prime) were used to visualize transferred proteins.

#### *In vitro* Assays

##### *In vitro* CAR Kinetic Activity Assay

*In vitro* CAR assays were performed in 95 µL of kinetic buffer was loaded into a 96 well plate. Substrates were dissolved in pure DMSO (Fisher Scientific) at 100 mM. Prior to spectrophotometric analysis of NADPH depletion, 5 µL of substrate was added to each reaction well, resulting in final composition of kinetic buffer composition of 100 mM HEPES pH 7.5 at 25°C, 0.5 mM NADPH (Santa Cruz Biotechnology, tetrasodium salt), 1 mM ATP (Sigma-Aldrich), 10 mM $MgCl_2$, 1.5 µM CAR. Oxidation of NADPH was measured at 25 ºC for triplicate samples using an Agilent BioTek Synergy H4 Hybrid Microplate Reader at a wavelength of 340 nm for 15 min. Spectral scans of substrates at and products 5 mM determined that there was no competing absorbance measurement at 340 nm (data not shown). All purified CAR preparations exhibited no appreciable oxidation of NADPH prior to addition of substrate. *In vitro* CAR activity was then analyzed to determine the maximum rate of NADPH oxidation, using 0 mM and 0.5 mM NADPH standards for calibration.

Benzoic acid, TPA, 4FBA and TPAL, mmTPA, and mm4FBA were sourced from Sigma-Aldrich at highest purity. pAMB, pXYL, and pAMBA were sourced from AmBeed at highest purity. MHET was sourced from Advanced Chemblocks at highest purity. MHE-4FBA was synthesized as part of this paper.

##### Endpoint Bioconversion of TPAL and 4FBA by cvTA

1.5 µM cvTA (dimer basis), 100 mM HEPES pH 7.5, 5% DMSO, 10 mM substrate (TPAL or 4FBA, Sigma-Aldrich), 400 µM PLP (TCI Chemicals), 40 mM *i*Pr-$NH_2$ (*i*Pr-$NH_2$, Sigma-Aldrich) were added to an Eppendorf tube. Each reaction was run at 100 µL scale in triplicate. The reaction was allowed to proceed for 24 h at 30 ºC with shaking at 250 rpm prior to quenching with 200 μL 2M HCl and 700 μL methanol. Samples were chilled at -80 ºC for 1 h prior to centrifugation to clear insoluble protein precipitate. Samples were then analyzed by HPLC-UV (see **HPLC Methods**)

**Endpoint Bioconversion of 5 mM TPA to TPAL by srCAR with Sequential Addition of Cofactor Regeneration Enzymes**

Bioconversion of TPA by CAR without cofactor regeneration was performed in 100 mM HEPES pH 7.5, 12 mM ATP, 12 mM NADPH, 20 mM $MgCl_2$, 5% v/v DMSO and 5 mM TPA. srCAR was added to a final concentration of 1.5 μM. For NADPH regeneration, 1.5 μM bmGDH, 1 mM $NADP^+$, and 50 mM glucose were added in place of 12 mM NADPH. For ATP regeneration, 0.42 mg/mL of PPK12, 1 mM AMP, and 15 mM SHMP were added in place of 12 mM ATP. Endpoint reactions were performed in triplicate at 100 μL and incubated at 30 ºC with shaking at 250 rpm. After 24 h, the reaction was quenched with 0.5 volumes of 2M HCl and 3.5 volumes of ultra-cold methanol. Samples were chilled at -80 ºC for 1 h prior to centrifugation to clear insoluble protein precipitate. Samples were then analyzed by HPLC-UV (see **HPLC Methods**).

**Timepoint Analysis - Bioconversion of 5 mM TPA, 4FBA, MHET, and mmTPA by srCAR with Cofactor Regeneration**

Bioconversion of TPA by CAR with cofactor regeneration was performed in 100 mM HEPES pH 7.5, 1 mM AMP, 1 mM $NADP^+$, 20 mM $MgCl_2$, 15 mM SHMP, 50 mM glucose, 5% v/v DMSO, 5 mM substrate (TPA, 4FBA MHET or mmTPA), 1.5 μM bmGDH (tetramer basis), 1.5 μM ecPPase (hexamer basis), 0.42 mg/mL PPK12 and 1.5 μM srCAR. Reactions were performed in triplicate at 300 μL and incubated at 30 ºC with shaking at 250 rpm. At each timepoint, for samples containing TPA, 4FBA and mmTPA as substrate, 35 μL was removed from the reaction and quenched in 0.5 volumes of 2M HCl and 3.5 volumes of ultra-cold methanol. For timepoints containing MHET as substrate, 35 μL of the reaction mix was quenched in 4 volumes of ultracold methanol. Samples were chilled at -80 ºC for 1 h prior to centrifugation to clear insoluble protein precipitate. Samples were then analyzed by HPLC-UV (see **HPLC Methods**).

**Endpoint Bioconversion of 5 mM and 10 mM TPA to pXYL by srCAR and cvTA with Cofactor Regeneration**

Bioconversion of TPA to pXYL by srCAR and cvTA with cofactor regeneration was performed in 100 mM HEPES pH 7.5, 1 mM AMP, 1 mM $NADP^+$, 20 mM $MgCl_2$, 15 mM SHMP, 50 mM glucose, 5% v/v DMSO, 5 or 10 mM TPA, 4x molar excess of *i*Pr-$NH_2$ (relative to TPA), 250 nM bmGDH (tetramer basis), 1.5 μM ecPPase (hexamer basis), 0.42 mg/mL PPK12, 1.5 μM srCAR and 1.5 μM cvTA. Reactions were performed in triplicate at 100 μL and incubated at 30 ºC with shaking at 250 rpm. After 24 h, the reaction was quenched in 0.5 volumes of 2M HCl and 3.5 volumes of ultra-cold methanol. Samples were chilled at -80 ºC for 1 h prior to centrifugation to clear insoluble protein precipitate. Samples were then analyzed by HPLC-UV (see **HPLC Methods**).

**HPLC Methods**

**HPLC Method 1: Detection of TPA, 4FBA, TPAL, pAMB and HMB**

TPA, 4FBA, TPAL, pAMB and HMB were quantified using reverse-phase high-performance liquid chromatography (RP-HPLC) with an Agilent 1260 Infinity with a Zorbax Eclipse Plus-C18 column with a guard column installed. To achieve separation of the mobile phase initially consisted of solvent A/B = 95%/5% (solvent A: water with 0.1% trifluoroacetic acid, solvent B: acetonitrile with 0.1% trifluoroacetic acid) and maintained for 25 min. A gradient elution (A/B) was performed from 25-27 min from 95/5 to 70/30, followed by a gradient from 27-30 min from 70/30 to 95/5, followed by isocratic hold from 30-32 min at 95/5 to equilibrate the column. A flow rate of 1 mL/min was used for all stages of the method and UV absorbance was quantified at 254 nm using commercially available standards for reference. The column temperature was maintained at 30 ºC.

**HPLC Method 2: Detection of MHET, MHE-4FBA mmTPA, and mm4FBA**

MHET, MHE-4FBA mmTPA, and mm4FBA were quantified using reverse-phase high-performance liquid chromatography (RP-HPLC) with an Agilent 1260 Infinity with a Zorbax Eclipse Plus-C18 column with a guard column installed. The mobile phase initially consisted of solvent A/B = 95%/5% (solvent A: water with 0.1% trifluoroacetic acid, solvent B: acetonitrile with 0.1% trifluoroacetic acid) and maintained for 5 min. A gradient elution (A/B) was performed from 5-10 min from 95/5 to 90/10, followed by a gradient from 10-15 min from 90/10 to 50/50, followed by gradient from 15-19 min from 50/50 to 95/5, followed by an isocratic elution from 19-20 min at 95/5 to equilibrate the column. A flow rate of 1 mL/min was used for all stages of the method and UV absorbance was quantified at 254 nm using commercially available standards for reference, except for 2-hydroxyethyl 4-formylbenzoate, which was chemically synthesized as part of this paper. The column temperature was held at 30 ºC.

**HPLC Method 3: Detection of pAMBA and pXYL**

Amines were derivatized with *ortho*-pthalaldehyde (OPA) and 3-mercaptopropionic acid and identified using reverse-phase high-performance liquid chromatography (RP-HPLC) with an Agilent 1260 Infinity with a Zorbax Eclipse Plus-C18 column with a guard column installed. The mobile phase solvents used were Solvent A: 10mM sodium phosphate, dibasic and 10mM sodium borate decahydrate in water, pH 8.2 and Solvent B: 45%/45%/10% (v/v/v) methanol/acetonitrile/water. Samples derivatization was automated during the pre-injection step using the HPLC manufacturer's protocol for OPA derivatization.[4] In brief, 2.5 μL borate buffer

(0.4N sodium tetraborate in water, pH 10.2) was mixed with 1 µL bioconversion sample. The sample was then reacted with 0.5 µL Derivatization Buffer (74.6 mM OPA, 94.2 mM 3-mercaptopropionic acid in Borate Buffer) and mixed in the sample needle. The sample was then diluted with 32 µL Injection Diluent (13.56 mM phosphoric acid in solvent A) and mixed prior to injection. The UV detector was set to 338 nm, 10 nm bandwidth, and reference wavelength 390 nm, 20 nm bandwidth, for analysis. The mobile phase initially consisted of solvent A/B = 98%/2% and maintained for 0.5 min. A gradient elution (A/B) was performed from 0.5-20 min from 98/2 to 43/57, followed by a gradient from 20-25 min from 43/57 to 0/100, followed by an isocratic hold from 25-35 min at 0/100, followed by a gradient elution from 35-39 min from 0/100 to 98/2, and a final hold at 98/2 from 39-42 min to equilibrate the column for the subsequent sample. A flow rate of 0.42 mL/min was used for all stages of the method. The injection needle was purged prior to derivatization of the next sample. The column temperature was maintained at 40 ºC.

#### Chemical Synthesis

##### Synthesis of 2-hydroxyethyl 4-formylbenzoate (MHE-4FBA)

A 2-dram vial was charged with 4-formylbenzoic acid (150 mg, 1 mmol), 2-bromoethanol (83 uL, 1.2 mmol), sodium bicarbonate (168 mg, 2 mmol) and dissolved in 3 mL of DMF. The vial was heated to 125 °C and stirred overnight. The vial was allowed to cool to room temperature before adding 2 mL of water. The organic layer was extracted with ethyl acetate (3 x 1 mL) and collected in a separate 20 mL scintillation vial. The organic solution was dried with magnesium sulfate, filtered, and concentrated via rotary evaporator. The crude mixture was purified by hexane/ethyl acetate in a 3:1 v/v ratio resulting in 104.5 mg of **2-hydroxyethyl 4-formylbenzoate** (54 % yield). $^{1}$H and $^{13}$C NMR spectra match those reported in the literature[5].

##### Synthesis monomethyl terephthalate (mmTPA) from TPA

Terephthalic acid (TPA) (8.31 g, 50 mmol) was refluxed in methanol (200 mL) for 30 min or until dissolved. Thionyl chloride (14.5 mL, 200 mmol) was then added dropwise to the solution and then refluxed overnight. The solution was allowed to cool to room temperature before concentrating *en vacuo*. The crude mixture was extracted with diethyl ether (2 x 50 mL) and then washed with 0.1M aqueous potassium hydroxide. The combined organic layers were dried with magnesium sulfate, filtered, and concentrated resulting in dimethyl terephthalate (DMT) and used without further purification. DMT (1.94 g, 10 mmol) was dissolved in methanol (33 mL) by stirring at 40 °C for 15 min. Potassium hydroxide (561 mg, 10 mmol) was then added before refluxing overnight. The solution was then cooled to room temperature and concentrated *en vacuo*. The crude was dissolved in water and extracted with ethyl acetate (3 x 10 mL). The aqueous layer was acidified with aqueous 1M HCl. The resulting precipitate was vacuum filtered and washed with water. The precipitate was left on the filter for 15 min to dry yielding monomethyl terephthalic acid (mmTPA) (1.51 g, 76% yield overall). $^1H$ NMR spectroscopy matched those previously reported in the literature[6,7].

#### Supplemental Tables

**Table S1.** Expression conditions and accession numbers for enzymes.

| Organism/Enzyme | Abbreviation | GenBank Accession Number | Cloning Method | Expression Condition | Genbank Plasmid Accession Number |
|---|---|---|---|---|---|
| *Valsa mali*/CAR | vmCAR | KUI69596.1 | Golden Gate | P1 | OP031622 |
| *Aspergillus fumigatus*/CAR | afuCAR | XP_748589.2 | Golden Gate | P1 | OP031615 |
| *Acremonium egyptiacum*/CAR | aeCAR | BBF25314.1 | Golden Gate | P1 | OP031613 |
| *Herbihabitans rhizosphaerae*/CAR | hrCAR | WP_130345796.1 | Golden Gate | P1 | OP031617 |
| *Streptomyces europaeiscabiei*/CAR | seCAR | WP_050362911.1 | Golden Gate | P2 | OP031620 |
| *Streptomyces griseus*/CAR | sgCAR | WP_012382217.1 | Golden Gate | P2 | OP031621 |
| *Streptomyces cyaneus*/CAR | scCAR | WP_128435011.1 | Golden Gate | P2 | OP031619 |
| *Mycobacteroides abscessus*/CAR | mabCAR | WP_005082584.1 | Golden Gate | P1 | OP031625 |
| *Streptomyces noursei*/CAR | snCAR | WP_102925360.1 | Golden Gate | P2 | OP031623 |
| *Mycolicibacterium smegmatis*/CAR | msCAR | AFP42026.1 | Golden Gate | P2 | OP031624 |
| *Cafeteria roenbergensis*/CAR | crCAR | KAA0160565.1 | Golden Gate | P2 | OP031628 |
| *Mycobacterium avium*/CAR | mavCAR | WP_003872682.1 | Gibson Assembly | P1 | OP031609 |
| *Trichoderma reesei*/CAR | trCAR | XP_006964071.1 | Gibson Assembly | P1 | OP031611 |
| *Neurospora crassa*/CAR | ncCAR | XP_955820.1 | Gibson Assembly | P1 | OP031610 |
| *Segniliparus rotundus*/CAR | srCAR | WP_013138593.1 | Gibson Assembly | P1 | OP031608 |
| *Nocardia iowensis*/CAR | niCAR | Q6RKB1.1 | Gibson Assembly | P1 | OP031607 |
| *Mycobacterium marinum*/CAR | mmCAR | WP_012393886.1 | Gibson Assembly | P3 | OP031612 |
| *Chromobacterium violaceum*/ω-transaminase | cvTA | WP_011135573.1 | Gibson Assembly | P1 | N/A |
| *Escherichia coli*/Inorganic pyrophosphatase | ecPPase | BAE78227.1 | Gibson Assembly | P1 | N/A |
| Unclassified *Erysipelotrichaceae*/Polyphosphate Kinase 2-III | PPK12 | HCY06753.1 | Gibson Assembly | P1 | N/A |
| *Bacillus megatarium*/Glucose dehydrogenase | bmGDH | WP_013055759.1 | Addgene 108954 | P1 | N/A |
| *Aspergillus flavus*/CAR | aflCAR | RAQ59689.1 | Golden Gate | Did not express under P1 or P2 | OP031614 |
| *Lasallia pustulata*/CAR | lpCAR | KAA6412420.1 | Golden Gate | Did not express under P1 or P2 | OP031618 |
| *Crucibulum laeave*/CAR | clCAR | TFK38857.1 | Golden Gate | Did not express under P1 or P2 | OP031616 |
| *Amphimedon queeslandica*/CAR | aqCAR | XP_019858162.1 | Golden Gate | Did not express under P1 or P2 | OP031626 |
| *Wallemia mellicola*/CAR | wmCAR | TIC21279.1 | Golden Gate | Did not express under P1 or P2 | OP031627 |

P1: LB with antibiotic, inoculation at OD600 0.1, incubated at 37 C until induction at OD600 of 0.5-0.8, incubation at 30°C for 5 h, followed by overnight expression to 18°C for 18 h. P2: Terrific broth with antibiotic, incubated at 37 C until induction at OD600 of 0.5-0.8, incubation at 30°C for 5 h, followed by overnight expression to 18°C for 18 h. P3: LB broth with antibiotic, incubated at 37 C until induction at OD600 of 0.5-0.8, followed by overnight expression at 18°C for 18 h.

**Table S2.** Oligonucleotides used in this study.

| Oligo Name | Sequence |
|---|---|
| pZE_BB_F | CTTGATGGGGGATCCCATGGTA |
| pZE_BB_R | GTGGTGATGATGGTGATGGCTGCTGCCCATGGTACCTTTCTCCTCTTTAATGAATTCG |
| pZE_niCAR_sfp_Ins_F | GCCATCACCATCATCACCACATGGCTGTGGACTCGC |
| pZE_niCAR_sfp_Ins_R | CCATGGGATCCCCCATCAAGAAGTTACAGCAGTTCTTCGTAGCTCAC |
| pZE_BB_sfp_F | GCGGCCGCTAATAAAAG |
| pZE_srCAR_Ins_F | GCCATCACCATCATCACCACATGACACAGTCTCACACTC |
| pZE_srCAR_Ins_R | TTTATTAGCGGCCGCTCATAATAAGCCCAACTGC |
| pZE_mabCAR_Ins_F | CACCATCATCACCACATGACCGAAACCATCTCAAC |
| pZE_mabCAR_Ins_R | TTTATTAGCGGCCGCTTAAACCAGGCCGAGC |
| pZE_mavCAR_Ins_F | CACCATCATCACCACATGAGCACGGCCACC |
| pZE_mavCAR_Ins_R | TTTATTAGCGGCCGCTTAGAGCAACCCCAGCAG |
| pZE_trCAR_Ins_F | CACCATCATCACCACATGCGTAGCTTCGTCAAAG |
| pZE_trCAR_Ins_R | TTTATTAGCGGCCGCTTACTGCCGGAGATAGCC |
| pZE_ncCAR_Ins_F | CACCATCATCACCACATGAGCCAACAGCAGAAC |
| pZE_ncCAR_Ins_R | TTTATTAGCGGCCGCTTAAAAATTCCACTGGCGACAC |
| pZE_mmCAR_Ins_F | GCCATCACCATCATCACCACATGTCCCCAATCACCC |
| pZE_mmCAR_Ins_R | CTCCTTTTATTAGCGGCCGCTTAAAGCAATCCAAGAAGGCG |
| pACYC_BB_F | AGCTTGATGGGGGATCC |
| pACYC_BB_R | CGAATTCGGATCCTGGC |

| | |
|---|---|
| pACYC_CvTA_ins_F | ACAGCCAGGATCCGAATTCGATGCAGAAGCAGCGTAC |
| pACYC_CvTA_ins_R | GGGGATCCCCCATCAAGCTTTCAGGCAAGTCCGCGAG |
| pACYC_ecPPase_ins_F | ACAGCCAGGATCCGAATTCGATGAGCTTACTCAACGTCC |
| pACYC_ecPPase_ins_R | AGGGGATCCCCCATCAAGCTTTATTTATTCTTTGCGCGCTC |

**Table S3.** DNA G-Blocks/Twist gene fragments for cloning in this study.
Start codons are underlined

| Oligo Name | Sequence |
|---|---|
| **PPK12** | ACAGCCAGGATCCGAATTCGATGATCAACATCTATAAGATCGATAAATTAAATAACTTTAACCTTAACAATCACAAAACGGACGA<br>TTACTCCCTGTGCAAGGATAAAGATACCGCATTAGAGCTGACACAGAAGAATATTCAAAAGATCTACGATTACCAGCAAAAACTT<br>TACGCAGAGAAAAAGGAAGGCTTGATCATCGCTTTTCAAGCGATGGATGCTGCGGGCAAGGACGGCACGATCCGTGAAGTCCTGA<br>AAGCCCTGGCGCCGCAAGGAGTTCATGAAAAACCGTTCAAGTCCCCATCAAGTACAGAGTTGGCCCATGATTATTTATGGCGCGTT<br>CACAATGCCGTTCCCGAAAAAGGGGAGATTACCATTTTCAACCGTTCTCATTATGAGGATGTCTTAATTGGCAAAGTTAAAGAACT<br>TTACAAGTTTCAGAACAAAGCAGACCGTATCGACGAGAATACTGTTGTCGATAATCGCTACGAAGATATTCGCAACTTCGAGAAGT<br>ATTTATACAACAACTCCGTGCGTATCATTAAGATCTTTCTTAACGTGTCCAAGAAGGAGCAAGCCGAACGCTTCTTGAGTCGCATT<br>GAAGAACCAGAAAAGAATTGGAAATTCAGTGACTCTGACTTCGAGGAACGCGTCTACTGGGATAAATACCAACAGGCTTTCGAAG<br>ATGCCATTAACGCAACTTCTACAAAGGACTGCCCGTGGTACGTTGTGCCGGCTGACCGTAAATGGTACATGCGCTATGTAGTCTCT<br>GAGATTGTGGTAAAAACCTTAGAAGAGATGAACCCGAAGTATCCAACGGTTACAAAAGAAACGTTGGAACGTTTTGAAGGCTACC<br>GCACGAAATTGCTGGAAGAATATAACTATGACCTTGACACAATTCGTCCAATCGAAAAGTGAAAGCTTGATGGGGGATCCCC |
| **srCAR** | ATGACACAGTCTCACACTCAAGGCCCTCAGGCGTCAGCAGCCCATTCCCGCTTGGCACGTCGTGCGGCAGAGCTCCTTGCGACCGA<br>TCCACAGGCGGCGGCAACGCTCCCTGACCCTGAGGTTGTCCGTCAAGCGACCCGTCCGGGCTTACGTCTCGCGGAGCGCGTCGACG<br>CGATCTTGAGTGGTTACGCCGATCGCCCTGCCCTTGGCCAGCGCTCTTTCCAGACTGTTAAAGATCCTATCACGGGACGTAGCAGT<br>GTAGAGTTATTGCCTACCTTTGACACGATTACGTACCGCGAGCTGCGTGAGCGTGCGACCGCCATCGCCAGCGATTTGGCCCACCA<br>CCCGCAGGCGCCGGCGAAGCCGGGTGATTTCCTCGCAAGTATTGGCTTCATTAGCGTCGACTATGTGGCTATTGATATTGCAGGCG<br>TTTTCGCGGGCCTCACGGCGGTGCCACTGCAAACTGGTGCAACCTTAGCAACGCTTACAGCGATTACGGCGGAAACCGCGCCGACC<br>TTATTCGCAGCGTCCATCGAGCATTTGCCTACTGCCGTAGACGCTGTCCTTGCTACGCCCAGTGTTCGCCGTCTTCTGGTATTTGACT<br>ATCGTGCAGGATCCGACGAGGACCGCGAGGCCGTCGAGGCAGCTAAGCGCAAGATTGCCGACGCCGGTTCCAGCGTGCTGGTTGA<br>CGTACTTGACGAAGTTATCGCCCGTGGCAAGAGTGCGCCAAAGGCTCCTCTCCCGCCAGCAACGGACGCAGGAGATGATAGCCTG<br>AGCTTGCTCATTTATACATCAGGCAGCACCGGTACTCCGAAGGGCGCGATGTATCCCGAGCGCAACGTCGCGCACTTCTGGGGCGG<br>AGTTTGGGCCGCCGCATTCGATGAGGACGCCGCACCCCCGGTACCGGCAATTAACATTACCTTCTTACCTCTCTCGCACGTTGCCTC<br>ACGTCTCTCTCTTATGCCTACCCTGGCGCGTGGCGGCCTGATGCACTTTGTTGCTAAGTCTGACCTGTCTACCTTGTTTGAGGACTTA<br>AAGCTGGCGCGTCCGACAAATCTGTTTCTTGTCCCACGTGTAGTCGAGATGCTCTATCAACACTACCAGTCCGAACTCGATCGTCG<br>CGGCGTACAAGACGGAACGCGTGAGGCGGAAGCGGTAAAGGATGACCTGCGCACGGGACTGCTGGGCGGGCGCATCCTCACTGC<br>CGGATTCGGTTCCGCCCCGTTGTCGGCGGAGCTGGCCGGTTTCATCGAGTCTTTACTTCAAATCCATCTGGTGGACGGATACGGCA<br>GCACCGAGGCCGGCCCGGTTTGGCGTGACGGTTATTTAGTAAAGCCGCCGGTGACGGATTACAAGCTTATTGACGTTCCCGAGCTT<br>GGTTATTTCTCTACAGACAGCCCTCACCCACGCGGTGAATTAGCGATTAAGACGCAGACCATCTTACCCGGATATTACAAACGTCC<br>GGAGACCACAGCTGAGGTGTTCGACGAGGACGGCTTTTATTTGACCGGTGACGTTGTGGCCCAAATCGGTCCCGAGCAATTCGCGT<br>ATGTGGACCGCCGCAAGAATGTTTTAAAGTTAAGTCAGGGTGAGTTCGTCACCTTAGCGAAGTTAGAGGCAGCATATTCGTCGTCC<br>CCTCTTGTACGTCAACTTTTCGTGTACGGCTCGAGTGAGCGCTCTTATCTGTTAGCCGTTATTGTCCCCACGCCTGACGCCTTGAAG<br>AAGTTTGGTGTCGGCGAGGCCGCCAAGGCCGCCTTAGGTGAGTCACTCCAGAAAATTGCGCGCGACGAGGGCCTGCAATCGTACG<br>AGGTTCCCCGCGACTTCATCATTGAGACAGATCCATTTACTGTAGAAAACGGCCTGTTAAGCGACGCACGTAAGAGCTTGCGTCCA<br>AAGCTCAAGGAGCACTATGGAGAGCGCCTCGAGGCGATGTACAAAGAGCTGGCTGATGGCCAGGCCAATGAGCTGCGCGATATTC<br>GTCGCGGCGTACAACAGCGCCCCACTCTGGAGACTGTGCGCCGTGCGGCGGCAGCAATGCTCGGTGCGAGCGCGGCTGAGATTAA<br>GCCAGACGCCCATTTTACGGACTTGGGCGGCGACTCGCTTTCCGCATTGACGTTTTCGAACTTCTTGCATGACTTGTTCGAGGTCGA<br>CGTGCCGGTGGGCGTAATCGTGTCGGCCGCGAATACTCTCGGTTCCGTCGCTGAACATATCGATGCGCAACTCGCAGGCGGCCGCG<br>CTCGTCCGACATTCGCCACCGTACATGGCAAAGGTTCAACAACCATTAAGGCCAGCGATTTGACGCTTGACAAGTTTATCGATGAA<br>CAGACCTTAGAAGCGGCTAAGCATCTGCCTAAACCAGCCGACCCGCCTCGCACCGTCTTGTTGACTGGGGCTAACGGGTGGTTGGG<br>CCGTTTCCTTGCCCTTGAATGGTTAGAACGCTTAGCCCCGGCCGGCGGTAAGTTAATCACGATCGTCCGCGGTAAGGACGCGGCGC<br>AAGCAAAGGCCCGCCTTGACGCCGCGTACGAGAGTGGGGATCCGAAGTTGGCGGGACACTACCAAGACTTAGCGGCAACGACGCT<br>GGAAGTATTGGCGGGAGATTTCTCAGAGCCCCGCTTAGGTTTGGACGAGGCAACCTGGAATCGTCTCGCAGATGAGGTTGACTTCA<br>TCTCGCATCCTGGCGCCCTTGTAAATCATGTTCTGCCTTACAATCAATTATTCGGCCCCAACGTAGCCGGCGTTGCGGAGATTATTA<br>AACTGGCAATCACCACGCGCATCAAGCCAGTGACGTACCTGAGTACGGTCGCGGTGGCTGCCGGGGTTGAGCCTTCAGCATTAGA<br>CGAAGATGGCGACATCCGCACGGTTTCAGCAGAGCGTAGTGTTGATGAGGGTTACGCCAATGGTTATGGCAACAGTAAGTGGGGT<br>GGTGAGGTGCTGTTACGCGAGGCCCACGACCGCACTGGCTTGCCTGTGCGCGTATTTCGTTCCGACATGATCCTGGCACACCAGAA<br>GTATACTGGCCAGGTAAATGCTACCGACCAGTTTACCCGTCTGGTACAATCGCTCCTGGCCACGGGTCTGGCCCCTAAATCGTTCT<br>ATGAGTTAGACGCGCAGGGTAATCGTCAACGTGCACATTACGACGGTATCCCTGTGGACTTCACAGCCGAGTCCATTACCACGTTG<br>GGTGGCGACGGGTTGGAAGGATATCGCTCATACAACGTGTTCAATCCTCATCGTGACGGCGTAGGCCTTGACGAATTCGTTGACTG<br>GCTCATCGAGGCAGGCCATCCGATCACGCGCATCGACGATTATGATCAATGGTTGAGTCGCTTCGAAACTAGTCTGCGTGGCTTAC<br>CGGAGAGTAAGCGCCAAGCGTCAGTCCTTCCGCTGCTTCACGCGTTCGCCCGCCCCGGTCCCGCGGTGGATGGTAGTCCGTTCCGC<br>AATACCGTGTTCCGTACGGACGTCCAGAAGGCCAAGATTGGCGCCGAACATGACATTCCTCACCTCGGTAAGGCGCTCGTTTTAAA<br>GTATGCGGACGACATCAAGCAGTTGGGCTTATTATGA |
| **trCAR** | ATGCGTAGCTTCGTCAAAGCGAACGTCGATTTCTCATCTGCCGAACGTAAGGAAGATTATATTCACAGCTTACCCGAATTGGTGGA<br>TTTTAACGCAGTTCAAAACCCAAACCATCTTCTTTGCATCCAGGCGCGCTCCAACGCCCCTTGGGTAAAAATCACGAATGCCCAAT<br>TCAAAGTCGCGATTGACCAATGTGCTACCTGGATTGCTGAGAATGTGAAATTACCGAAAGCGCGTACAAAGCATGACCTGACTGG<br>CCGCCTGCCGGTGGCACTGCTGATGGAATCAGACTTTGGCCTGCTTGTCCATCAATTCGCGCTGGTGAGTATGGGGATTCCGCCAC<br>TGGTGCTGAGTGCCCGGTTGTCCCCGGAAGCTATCTTTCATTTACTTCGCAGTACCGAGGCGTCCTCTCTGATCGTTAGCCAGCGTG<br>TTGCAATGATCACCAAAGGGGCATTCGGGAATGTCAAGACCTCCGACTTCCATGTGGCACAGCCGTACAGTACGTTTTGTAATGTG<br>CCGGCGGACAAAAGCGTTCGCAAACAGAGCGTGTATCCAGACAATATTGATGCCAATATCGTGTTGCTGCATAGTTCGGGCACCA<br>CAGGCTTACCGAAACCGATCGCCCTGAGCCACCGGCAGCTGATGTTCAGCGTCAGCCACGGGGATTTTGAAACCGAAGAAGAAGC<br>GCAGGGCATTGTTATCAGTACCCTTCCGCTTTTCCACGGATTTGGCCTGCTCGCGCCGGGTCTGAGCATGGCAATCGGTAAGACCG<br>TGTGTTTTCCCGCGAGTGATGAGGTTCCGGATGCCCAGAGCATTGTAGATCTTATCAACATGTCTGGCGCAACCGGCATGCTGACG<br>GTGCCCTTCTTGCTTGAGAACATGGCCGCATTACCGAACGGCACTGGTTTACGCGCACTGGCCAAGCTGGATTTTGTGGGGACTGG<br>TGGCTCCGCGCTGAGTGCAGATTTTGGCGTCTCAGCCTCTGCGGCGGGCGTCAAACTGCTTAACTTGTACGGCACCACAGAGACCG<br>GACCTTTGACCAAAACGTTCGCCCCGAAATCGGGCTATGATTGGAAGTACTTTCGCTTACGTCAGGATATGCTGTTTAAGGTGACA |

| | |
|---|---|
| | GAGCTCCCACCAGTCGATGGTGAAAAGCGTTTCCGGCTGACGGTCTTTCCGTTCGGTGCAGATAAACCTTTTGAAATTGCTGATCA<br>GCTCATCCGTTCCGAGAAGTTCCCGGAGACCGACTTCGCCGCCGTCGGGCGCGACGATGATGTGGTAGTGTTGGCGACGGGCGAA<br>AAAGTGAACCCACTGCTGCTGGAAACCGCATTGACCGACTCAGGACTGGTCAAGAGCGCAATCGTATTTGGGGAAAATCAATTCC<br>AAATCGGCGTAGTGGTGGAGCCTGCGACGCCCTTAAATCCCGATCAGAAAGAAGAATTTCGCAAAAAAATCTGGCCCATCATCGT<br>CCGTGTTGGCGAACGCATGGATACGACCGCACGCATTTATTCGCCAAACGCGGTTATTGTTGTTCCAAGCTCAGTGACCATCCCGC<br>GCACGGATAAAGGGTCGATCGCTCGCAAAGAAGTTTTTCAGCTGCTTGAGAAGGAGATTTCTCAGGTCTACGAGGATTTAGAAAAT<br>GGTTCGATTGAAGAAACACCCTTAGACTATGACAAACTGGAACAAGAACTGAAAGGTCTTATCCAGAAGCGCCTTAAACTGCGCG<br>TTCACCCGGGTAAGTGGACCGTAGACGATAATCTTTTTCATCTGGGACTCGATTCTCTGCAAGCGACCACGCTGCGTCGTATTTTAC<br>TTAGCGCCGCCTCGAAGACCCCGCCGGATGTTATTGGCAAAGATTTTATCTATGTAAACCCAAGTGTGAAAGCAATTGCTAACGCA<br>CTGCGCCCTGCGAATGGTCCAATTGGCACGGAGTCTGCCTCTGTTGCCCAGGAAGTGGATGATTATGCACAGCAGTACAGCATTAA<br>AGGCTTTGAAGTGCAAGACATCGTACCAAAGGCATCCCCGAAACTTATCCGCGGCGCGGTGGTTCTGCTCACAGGGTCCTCAGGCG<br>GCCTCGGTAGTCATGCCCTGGGCAAGCTGGCCGAGTCAACTCAGGTTGCCAAAATTGTCTGCTTGCAGCGGAAACGCCCGGGCAC<br>AGTAATCAACCCTATTCCCGGTGCGGCAAAAGTAGATCGCGCCTCCATTGAGGCAAAGGGCATTAAACTGACCGACGATCAGTGG<br>GCAAAAATTACGGCGCTGGAAATTGATCCGACGATTGATAATCTGGGGCTGCCGGCGATGGTGATGGGTATGGTTTCTAAGACCGT<br>CACGCACATCTTACACGCAGCTTGGCCGATGGACTTTCACATGCGTCTTCCATCATTTGGTTACCAGTTCAGCTATTTAAAGAACCT<br>CCTGCGTATTGCCGTCCAGGCCCCGCAGAAAGTTCGCTTCCTGTTTGTATCTTCAATTAGCGCACTGGCAAAGCTGGGTCTGATTAC<br>CCCAGGCCGCCCGATCCCAGAAGAACCCCTCGATGTCGAAAGCGCGGCTTGCGGCATTGGCTATGCTGATGCGAAACTTGTCTGCG<br>AGAAAATCCTGGAAGAAGCAGCATCGCTCTATAACAGTAATGTTGAGGTGGTGATCGCCCGTTGCGGCCAGCTGAGCGGCGCACG<br>CAAGACCGGTGCGTGGAACGTCTCCGAACAGATCCCGATGCTTATCCGCACGAGCCAAGGTTTAGGAATTCTGCCGATCCTTGAAG<br>GCACCGTGTCCTGGATCCCGGTCGATGACGCAGCGGCCACCGTGGCAGAGTTACTCTTTGCCCCCGATGCGCCCGGTCTTGTGACG<br>CATGTTGAAAATCCGGTCCGTCAGTCATGGAGCGAAGTATTCCAAATCATCGGCAACGAACTGCGTATTACCAAAACACTGAGCTT<br>CGACGATTGGCTCGGCGAAGTGACATCAACCGCGGAACGTGATGTGGAGGATTACCCGGTGCGCAAACTGTATGAATTCTTTAAA<br>CTGTACTTCCGTATTGCCTCCTCAGGTGCGGTAGTTATGGGTACGGATATGAGCCGCAAAAATTCAGCCACGCTGCGCTGTCTTAA<br>AGCGCTCGACCGCGGAACTATTGCGGGCTATGTGCGCTATTGGCGCAGCGTCGGCTATCTCCGGCAGTAA |
| **ncCAR** | ATGAGCCAACAGCAGAACCCGCCATATGGACGTCGCCTTATTTTGGACATTATCAAGGAACGTGCCTTAAATGAACCGAATCGCG<br>AATGGGTCAGTGTGCCCCGCTCTAGCGATCCGAAAGATGGTTGGAAAATTCTGACCTACTTGGACGCGTATAATGGAATTAACCGT<br>GTAGCGCATAAACTGACCCAAGTATGCGGCGCTGCAGCACCGGGGTCATTCCCCACCGTCGCGTACATCGGACCGAACGATGTGC<br>GTTATCTTGTATTCGCACTGGGCGCCGTCAAGGCGGGCTACAAAGCCCTGTTCATTTCAACCCGGAATAGTGCAGAAGCCCAGGTT<br>AACTTATTCGAATTGACTAACTGTAACGTCCTGGTGTTCGACCAAAGTTACAAGGCGACTGTGCAACCGTGGTTACACGAGCGTGA<br>AATGACCGCTATTCTGGCGTTGCCGGCCGATGAATGGTTCCCGGCGGATCAGGAAGACTTTCCTTATAACAAAACCTTTGAAGAGG<br>CCGAATGGGATCCGCTTATGGTACTGCATACCAGTGGGTCAACAGGCTTTCCTAAGCCGATTGTTGCTCGTCAGGGCATGCTGGCG<br>GTCGCCGACCAGTTCCATAACCTGCCTCCACGCGAAGATGGTAAGTTAATGTGGATCGTTGAGATGTCGAAACGCGCCAAGCGCTT<br>AATGCATCCGATGCCACTGTTCCACGCCGCCGGTATGTATATCAGCATGTTAATGATTCATTATTGGGATACCCCGGGCGCCTTAG<br>GCATTGGCGAACGCCCGCTGAGTAGCGATTTGGTTCTGGACTATATTGAATACGCGGATGTGGAAGGTATGATCTTGCCGCCCGCA<br>ATTCTCGAGGAACTGAGCCGCGACGAGAAGGCGATCCAGTCACTTCAGAAGCTGAACTTTGTCTCCTTTGGTGGCGGTAATTTAGC<br>GCCTGAGGCAGGTGACCGCCTGGTGGAGAACAACGTGACTCTGTGTAATCTTATCTCGGCCACGGAATTCACCCCGTTCCCGTTCT<br>ACTGGCAATATGATCAGAAACTTTGGCGCTATTTTAATTTTGATACTGATCTGTTTGGTATTGACTGGCGGTTACACGACGGTGAGA<br>GTACGTACGAACAAGTCATTGTGCGGAAAGATAAGCATCCGGGCTTACAAGGTTTCTTCTATACCTTTCCAGACTCATCGGAATAC<br>AGTACGAAAGATCTGTATAAACGCCATCCGACGCACGAGGATTTTTGGATTTATCAGGGACGTGCTGATAACATTATTGTTTTCTCC<br>AACGGAGAAAAACTCAACCCTATTACCATTGAAGAAACGTTACAAGGACACCCGAAAGTTATGGGCGCAGTTGTCGTAGGTACGA<br>ACCGTTTCCAACCTGCCCTGATTATTGAACCGGTTGAACACCCGGAAACAGAAGAGGGTCGTAAAGCTCTTCTGGATGAAATTTGG<br>CCTACCGTGGTACGCGTGAATAAGGAGACCGTTGCCCATGGCCAGATCGGCCGCCAGTATATGGCGCTGTCCACCCCCGGCAAAC<br>CGTTTCTCCGTGCCGGTAAGGGCACTGTGCTGCGTCCCGGCACAATTAACATGTATAAAGCAGAAATTGATAAAATTTATGAGGAT<br>GCGGAAAAGGGCGTTGCCACGGATGAGGTGCCAAAGCTGGATCTGTCGAGCAGTGACGCGTTAATTGTTTCCATCGAAAAATTGT<br>TCGAGACGTCACTGAATGCTCCGAAACTGGAGGCCGATACAGACTTTTTCACGGCTGGCGTTGATTCGATGCAGGTAATCACCGCG<br>TCTCGCCTGATTCGCGCGGGGCTGGCAGCTGCGGGCGTTAATATCGAAGCATCTGCGCTGGCGACTCGTGTGATTTATGGAAATCC<br>CACCCCAAAACGTCTTGCGGATTATTTGCTGTCGATCGTGAATAAAGACTCAAACCAGGGCACGTTGGACAATGAACATCATGTGA<br>TGGAAGCGCTGGTTGAGAAATACACGCGCGACTTGCCTACGCCGAAACAAAACAAGCCGGCGCCTGCCGACGAAGGTCAAGTTGT<br>GGTCATCACTGGTACCACCGGCGGGATCGGTTCTTATCTGATCGACATTTGCAGCAGCTCCAGTCGCGTTAGCAAGATTATTTGTCT<br>GAACCGCTCGGAAGACGGTAAAGCGCGCCAAACCGCAAGCTCCTCCGGTCGCGGCCTGTCGACGGATTTCTCCAAGTGTGAATTTT<br>ACCATGCGGACATGAGTCGGGCGGACCTTGGTCTGGGCCCGGAAGTCTATTCTCGCCTCCTGTCGGAAGTCGATCGGGTGATCCAT<br>AATCAGTGGCCCGTCAACTTCAATATTGCCGTGGAATCTTTCGAGCCGCACATCCGTGGTTGTCGCAATTTGGTGGATTTCTCCTAC<br>AAAGCAGATAAAAACGTCCCGATCGTATTTGTTAGTTCTATCGGCACGGTCGACCGCTGGCATGACGAAGACCGTATTGTTCCCGA<br>AGCTTCACTGGATGACCTGTCCTTGGCCGCCGGCGGTTACGGCCAGAGCAAATTAGTAAGCAGCTTGATCTTTGATAAAGCAGCTG<br>AAGTTAGCGGCGTCCCAACCGAAGTGGTTCGCGTTGGTCAGGTTGCTGGTCCTTCAAGCGAAAAAGGGTACTGGAATAAGCAGGA<br>GTGGCTGCCAAGCATCGTAGCGTCGTCCGCTTATTTGGGTGTTCTTCCTGACTCCCTGGGACAAATGACCACCATCGATTGGACTCC<br>AATCGAAGCCATTGCGAAATTGCTGCTGGAAGTTAGTGGTGTTATTGATAATGTGCCGCTGGATAAAATTAACGGATACTTCCACG<br>GTGTTAACCCTGAGCGGACCAGCTGGTCCGCTCTGGCTCCAGCGGTGCAGGAGTATTACGGTGACCGTATTCAGAAGATCGTTCCG<br>CTGGATGAGTGGCTCGAGGCGCTGGAAAAATCCCAGGAAAAGGCCGAAGATGTCACCCGCAATCCGGGTATTAAGCTGATTGATA<br>CTTATCGTACGTGGTCAGAAGGCTACAAAAAGGGTACGAAGTTTGTACCACTGGATATGACTCGCACTAAAGAATACTCGAAAAC<br>GATGCGTGAAATGCATGCAGTCACGCCTGAGCTGATGAAGAATTGGTGTCGCCAGTGGAATTTTTAA |
| **mavCAR** | ATGAGCACGGCCACCCATGATGAACGCTTAGACCGCCGCGTGCACGAGCTGATTGCAACTGACCCTCAGTTTGCGGCGGCTCAACC<br>TGATCCCGCAATTACGGCTGCTCTTGAGCAACCTGGTTTGCGTTTACCACAGATCATTCGTACAGTCCTGGACGGATATGCCGACC<br>GTCCGGCATTGGGACAGCGCGTGGTCGAGTTCGTTACCGACGCTAAAACCGGTCGTACTAGCGCTCAACTCCTGCCGCGCTTTGAG<br>ACCATCACATACGGAGAGGTGGCACAACGTGTTTCAGCCTTAGGTCGTGCACTGAGCGACGATGCGGTTCACCCGGGCGATCGTGT<br>TTGTGTGTTAGGGTTCAATAGCGTTGATTATGCAACTATCGACATGGCGCTCGGTGCAATCGGGGCGGTGAGCGTTCCACTCCAAA<br>CGTCAGCTGCCATTTCATCATTGCAGCCCATCGTAGCTGAGACTGAGCCAACACTTATCGCCAGCTCTGTGAATCAACTGTCCGAT<br>GCGGTCCAACTGATCACTGGCGCTGAGCAAGCCCCGACGCGCCTCGTCGTGTTCGACTACCACCCACAAGTGGACGACCAACGTG<br>AAGCAGTTCAAGACGCAGCTGCCCGTTTGTCAGGTACAGGTGTCGCGGTGCAAACATTGGCGGAGCTTTTAGAACGCGGTAAGGA<br>CCTTCCAGCCGTTGCTGAGCCACCCGCAGATGAGGATTCTTTGGCGTTATTGATCTACACTTCCGGTTCGACGGGTGCCCCCAAAG<br>GCGCGATGTACCCTCAGTCTAACGTGGGGAAGATGTGGCGTCGTGGTTCAAAGAACTGGTTTGGGGAGTCCGCAGCGTCTATTACC<br>TTAAACTTTATGCCCATGTCCCACGTGATGGGGCGCTCCATTCTGTACGGAACTTTAGGAAATGGCGGGACCGCCTACTTTGCAGC<br>CCGTTCGGATTTAAGCACCCTGTTAGAAGACCTGGAATTAGTACGCCCTACGGAGCTTAATTTTGTACCTCGTATCTGGGAGACTTT<br>ATATGGCGAGTTCCAACGCCAAGTGGAGCGTCGCCTTTCAGAGGCCGGGGACGCCGGCGAGCGTCGCGCGGTTGAAGCCGAAGTA<br>TTGGCCGAACAACGTCAATACTTGTTGGGCGGCCGCTTCACGTTCGCTATGACGGGATCGGCGCCGATTAGCCCCGAGCTGCGTAA<br>CTGGGTAGAGAGTCTGTTGGAGATGCACTTGATGGATGGTTACGGCAGTACTGAGGCAGGTATGGTGCTCTTTGACGGAGAAATTC<br>AGCGTCCGCCTGTCGTGGACTACAAGCTGGTAGATGTACCTGACTTAGGCTACTTTTCCACTGACCGTCCGCACCCGCGCGGTGAA<br>TTACTGTTACGTACTGAGAACATGTTTCCAGGCTATTATAAGCGTGCAGAAACGACTGCTGGGGTTTTCGACGAGGACGGTATTA<br>CCGCACTGGCGACGTGTTCGCCGAGATCGCGCCCGACCGCTTGGTGTATGTCGATCGCCGCAATAATGTGTTGAAGCTGGCGCAGG<br>GTGAGTTCGTGACGTTAGCCAAGCTCGAGGCGGTATTCGGGAACAGCCCTCTGATTCGTCAGATCTACGTCTACGGCAATTCGGCC |

| | |
|---|---|
| | CAGCCATATCTTCTTGCAGTTGTTGTTCCCACAGAAGAGGCACTGGCGAGTGGGGACCCAGAGACCTTAAAGCCAAAGATCGCCG<br>ACTCACTCCAGCAAGTAGCGAAAGAGGCCGGATTGCAATCCTATGAGGTGCCCCGTGACTTTATTATCGAAACGACACCGTTTTCC<br>TTAGAGAACGGGCTGCTTACAGGCATCCGTAAGTTGGCCTGGCCAAAACTCAAGCAACATTATGGTGAGCGCTTAGAGCAAATGT<br>ATGCAGATCTGGCGGCGGGGCAAGCAGATGAGTTAGCAGAACTCCGTCGCAATGGCGCCCAAGCGCCTGTTCTGCAAACTGTGTC<br>CCGCGCAGCAGGCGCAATGTTGGGGTCAGCGGCATCTGACCTGAGTCCCGACGCACATTTTACGGATTTAGGCGGCGATTCGCTCA<br>GCGCACTCACGTTTGGCAATTTGCTGCGTGAGATCTTTGATGTCGACGTGCCAGTAGGTGTAATTGTCAGTCCAGCGAATGATCTTG<br>CAGCCATCGCGTCGTATATTGAGGCTGAACGCCAGGGTTCGAAACGCCCTACCTTCGCGAGCGTTCATGGGCGTGACGCAACCGTT<br>GTGCGTGCGGCTGACCTGACTCTGGACAAGTTTCTTGATGCTGATACCCTTGCAAGTGCGCCCAATTTACCAAAACCAGCTACCGA<br>AGTGCGCACAGTCCTGTTAACCGGCGCGACAGGCTTCTTAGGTCGCTATCTGGCCCTTGAGTGGCTGGAACGCATGGACATGGTCG<br>ATGGAAAGGTCATTGCCTTAGTTCGCGCACGCAGCGACGAGGAAGCCCGTGCGCGTCTCGACAAGACGTTTGACTCGGGTGACCC<br>GAAGCTTCTTGCCCACTACCAACAATTAGCCGCCGACCACTTGGAAGTCATTGCAGGTGATAAGGGCGAGGCGAACCTTGGGCTG<br>CGTCAAGATGTATGGCAGCGTCTGGCCGATACAGTTGATGTCATCGTTGACCCGGCAGCATTAGTGAATCACGTGCTTCCTTATAG<br>TGAGTTATTTGGACCGAATGCCTTAGGCACGGCGGAATTAATTCGTTTAGCACTTACTAGCAAGCAGAAACCGTATACTTACGTCA<br>GTACCATCGGCGTCGGTGATCAAATTGAGCCTGGAAAATTCGTCGAGAATGCTGATATCCGTCAAATGTCCGCAACACGTGCGATC<br>AACGATTCTTATGCGAACGGTTACGGTAACAGTAAATGGGCTGGAGAGGTCCTCTTGCGTGAGGCTCATGATCTGTGCGGTCTGCC<br>CGTCGCGGTGTTTCGTTGTGATATGATTCTGGCCGACACTACATATGCCGGTCAATTAAATCTGCCAGATATGTTCACGCGCCTTAT<br>GCTGTCTTTAGTTGCCACGGGAATCGCACCAGGTAGCTTTTACGAACTGGACGCAGATGGCAATCGCCAACGTGCGCACTACGATG<br>GGTTGCCGGTCGAGTTCATCGCAGCGGCCATTAGTACCCTGGGTTCGCAGATCACAGACAGCGATACCGGCTTCCAAACATACCAC<br>GTAATGAATCCTTACGACGACGGGATTGGTCTTGACGAATATGTCGACTGGTTAGTGGACGCGGGATACAGCATTGAGCGCATTGC<br>AGATTATTCTGAATGGCTTCGTCGTTTTGAAACCAGTTTACGTGCACTGCCGGATCGCCAACGCCAGTATTCACTTCTTCCCTTACT<br>GCACAACTACCGTACGCCTGAGAAGCCTATTAACGGCAGCATTGCACCTACCGACGTTTTCCGCGCTGCCGTCCAGGAAGCGAAA<br>ATCGGTCCGGATAAGGACATCCCACATGTGAGCCCGCCGGTCATTGTCAAGTACATTACGGATTTGCAACTGCTGGGGTTGCTCTA<br>A |
| **mabCAR** | ATGACCGAAACCATCTCAACTGCAGCCGTACCAACAACGGATTTAGAAGAGCAGGTAAAACGCCGTATTGAGCAAGTTGTTAGTA<br>ACGATCCGCAGTTGGCCGCATTACTTCCCGAGGATAGCGTCACGGAAGCGGTTAACGAGCCGGATCTTCCGTTGGTTGAGGTGATT<br>CGTCGTCTGCTGGAAGGGTATGGTGACCGTCCTGCGTTAGGGCAACGCGCATTCGAATTTGTCACCGGTGACGACGGTGCGACAGT<br>TATTGCATTAAAACCAGAATACACGACGGTTAGCTACCGCGAACTGTGGGAGCGCGCCGAAGCTATTGCTGCCGCATGGCACGAG<br>CAGGGCATCCGTGATGGGGATTTCGTAGCCCAGCTCGGATTCACAAGCACGGACTTTGCCAGCCTGGACGTAGCGGGACTTCGCCT<br>GGGTACAGTGTCTGTGCCTTTACAAACGGGTGCGAGTCTGCAGCAACGTAACGCTATTCTCGAGGAAACGCGTCCAGCCGTTTTCG<br>CTGCTTCCATTGAATATCTTGATGCTGCTGTCGACTCAGTTCTGGCTACCCCCAGCGTGCGCCTCTTAAGTGTCTTTGACTACCATGC<br>CGAGGTCGACAGCCAGCGCGAGGCGCTTGAGGCAGTACGTGCCCGCTTAGAGAGCGCGGGTCGCACTATCGTGGTAGAAGCATTG<br>GCTGAAGCTCTGGCTCGCGGACGTGACCTTCCCGCCGCCCCGTTACCAAGTGCTGACCCAGACGCGTTACGTCTCCTCATCTACAC<br>CTCAGGTTCTACCGGAACCCCTAAGGGTGCTATGTACCCGCAATGGTTGGTCGCCAACTTGTGGCAAAAGAAATGGTTAACCGATG<br>ATGTTATCCCGTCGATTGGCGTGAATTTTATGCCAATGTCTCACTTGGCGGGACGTTTAACCCTTATGGGCACCCTGTCTGGCGGCG<br>GTACCGCCTATTATATCGCGAGCTCTGACTTGTCTACGTTCTTCGAAGACATCGCCCTCATCCGCCCTAGCGAGGTCCTCTTTGTCC<br>CACGCGTCGTGGAGATGGTATTCCAACGTTTTCAAGCCGAATTAGATCGTTCACTGGCACCGGGCGAGAGTAATTCGGAGATTGCT<br>GAACGTATTAAGGTGCGCATTCGCGAGCAAGATTTCGGCGGGCGCGTCCTGTCAGCAGGGAGCGGGTCAGCGCCACTGTCACCTG<br>AAATGACGGAGTTCATGGAGAGCTTACTCCAAGTCCCTCTGCGCGACGGGTATGGGTCTACTGAAGCGGGCGGCGTTTGGCGTGA<br>CGGGGTGCTTCAACGTCCGCCGGTCACAGACTATAAACTGGTTGACGTGCCCGAGTTGGGCTATTTCACCACGGATAGCCCACATC<br>CGCGCGGCGAACTTCGCTTGAAATCTGAAACAATGTTCCCAGGCTATTACAAGCGCCCGGAGACCACGGCGGACGTTTTCGACGA<br>CGAAGGGTACTATAAGACCGGTGACGTAGTTGCAGAGCTCGGGCCTGACCATCTGAAATATCTTGACCGTGTTAAGAATGTGCTGA<br>AGCTTGCCCAGGGTGAGTTCGTTGCCGTATCCAAACTGGAAGCGGCATACACGGGAAGTCCGCTTGTCCGTCAAATCTTTGTGTAT<br>GGGAACTCCGAGCGTTCCTTTCTTCTGGCTGTAGTGGTGCCCACGCCAGAGGTATTGGAGCGCTACGCCGACAGCCCCGACGCACT<br>GAAGCCCCTGATTCAAGACTCCCTTCAACAAGTGGCGAAGGACGCGGAGCTGCAATCGTATGAGATTCCACGTGACTTTATTGTGG<br>AAACGGTACCTTTTACGGTCGAGTCGGGCCTGCTGTCAGATGCTCGCAAGTTGTTACGCCCGAAGCTGAAGGACCACTATGGTGAG<br>CGTCTTGAGGCCTTATACGCCGAGTTAGCCGAAAGCCAAAACGAACGTCTCCGTCAACTCGCACGTGAGGCCGCGACGCGTCCCGT<br>CTTAGAGACAGTAACGGATGCTGCTGCGGCGCTGCTTGGAGCTAGCAGCTCAGATTTAGCACCAGATGTTCGCTTTATTGACCTCG<br>GCGGCGACAGTTTATCGGCATTGTCGTACTCGGAGCTGTTACGCGACATCTTCGAGGTTGACGTCCCTGTAGGAGTGATCAATTCG<br>GTGGCGAATGACCTTGCGGCGATTGCTCGTCATATTGAGGCCCAGCGCACGGGCGCCGCTACTCAACCCACTTTCGCTAGCGTGCA<br>CGGAAAGGATGCGACCGTTATCACGGCCGGAGAATTAACTCTGGATAAGTTTCTCGACGAGTCTTTGCTGAAGGCGGCGAAGGAC<br>GTGCAACCAGCCACGGCGGATGTGAAGACCGTGCTGGTCACGGGTGGCAATGGTTGGCTGGGTCGCTGGTTAGTCTTGGATTGGTT<br>GGAACGCCTGGCACCCAATGGCGGAAAAGTATATGCACTCATCCGCGGAGCAGATGCAGAGGCCGCCCGCGCGCGTCTGGACGCA<br>GTTTACGAGAGTGGTGATCCCAAGCTGAGCGCCCATTACCGTCAACTCGCGCAGCAAAGCTTAGAAGTTATCGCGGGTGACTTCGG<br>CGATCAAGACCTTGGTTTATCACAAGAGGTGTGGCAGAAATTAGCAAAAGATGTCGACTTAATCGTGCATTCTGGCGCGTTGGTGA<br>ACCACGTACTGCCGTACTCGCAACTGTTTGGGCCCAATGTGGCGGGCACTGCAGAGATTATTAAGTTAGCGATCAGTGAACGTTTA<br>AAGCCGGTTACGTACTTGTCTACCGTTGGTATCGCAGACCAGATTCCCGTTACCGAGTTTGAGGAAGATTCAGACGTTCGCGTCAT<br>GTCAGCCGAGCGCCAAATTAATGACGGCTATGCCAATGGTTACGGTAACTCCAAGTGGGCAGGTGAGGTACTTCTCCGTGAGGCA<br>CACGACCTGGCAGGTTTACCAGTGCGTGTCTTCCGCTCGGACATGATTCTCGCCCACTCAGATTATCATGGGCAATTAAACGTAAC<br>GGACGTTTTCACGCGTAGCATCCAGTCGCTGCTTCTTACAGGCGTTGCGCCAGCCAGCTTCTACGAACTTGATGCCGACGGTAATC<br>GTCAACGCGCGCACTACGACGGCGTGCCAGGCGATTTTACAGCAGCCAGTATCACCGCGATTGGTGGAGTAAACGTCGTGGACGG<br>CTATCGTTCTTTTGACGTCTTTAACCCACATCACGACGGAGTGAGTATGGATACCTTTGTGGACTGGCTCATTGACGCAGGGTATAA<br>GATCGCTCGCATCGATGACTACGACCAGTGGCTGGCTCGTTTCGAATTGGCATTGAAGGGTCTTCCGGAACAACAGCGTCAACAAT<br>CGGTCCTTCCTCTCTTAAAGATGTATGAGAAGCCACAACCGGCTATTGATGGGTCGGCCCTTCCGACCGCGGAGTTCTCTCGTGCA<br>GTACACGAGGCAAAGGTTGGGGACTCAGGGGAGATTCCGCACGTAACAAAGGAGCTGATTTTAAAATACGCCTCAGACATCCAAC<br>TGCTCGGCCTGGTTTAA |
| **cvTA** | ATGCAGAAGCAGCGTACAACATCGCAATGGCGCGAACTTGACGCCGCTCATCACCTGCATCCCTTCACCGATACCGCCTCCCTTAA<br>CCAGGCCGGCGCGCGCGTGATGACACGTGGAGAAGGGGTGTATTTGTGGGACTCGGAGGGAAATAAAATCATCGACGGTATGGCT<br>GGATTATGGTGTGTGAACGTTGGCTACGGTCGTAAGGACTTTGCCGAAGCGGCCCGTCGTCAGATGGAAGAATTACCGTTCTACAA<br>TACTTTTTTCAAAACAACCCATCCTGCGGTCGTAGAGTTATCTTCATTATTGGCGGAAGTCACTCCAGCAGGGTTTGACCGCGTGTT<br>TTATACAAATAGTGGATCAGAATCGGTTGACACAATGATCCGTATGGTCCGTCGTTACTGGGACGTCCAAGGCAAACCGGAGAAG<br>AAGACGTTAATCGGCCGCTGGAATGGTTATCACGGTTCGACCATTGGAGGTGCATCTCTTGGGGGCATGAAGTATATGCATGAGCA<br>GGGTGATTTGCCTATCCCTGGCATGGCGCACATCGAACAACCGTGGTGGTATAAGCACGGTAAAGACATGACGCCGGACGAGTTT<br>GGAGTTGTCGCTGCGCGTTGGTTGGAAGAGAAGATCCTGGAAATTGGGGCGGACAAGGTAGCCGCCTTCGTAGGAGAACCAATCC<br>AAGGTGCCGGGGGAGTGATCGTCCCGCCAGCTACCTATTGGCCCGAGATCGAGCGCATTTGCCGTAAATATGACGTATTGCTGGTT<br>GCAGATGAGGTAATTTGTGGCTTCGGGCGCACCGGGGAGTGGTTCGGGCACCAACATTTCGGTTTTCAGCCGGACTTATTTACGGC<br>GGCGAAGGGTTTAAGCTCAGGTTATTTACCGATTGGGGCTGTGTTTGTGGGCAAGCGTGTTGCCGAAGGCTTAATCGCGGGAGGCG<br>ACTTTAATCACGGATTCACATACTCTGGACACCCGGTTTGTGCCGCAGTAGCTCACGCGAATGTAGCCGCATTACGTGACGAGGGA<br>ATCGTCCAGCGTGTGAAGGACGATATCGGCCCTTATATGCAGAAGCGCTGGCGCGAGACTTTTTCACGTTTTGAGCACGTAGACGA<br>TGTGCGTGGCGTAGGCATGGTACAGGCCTTTACCTTAGTCAAAAATAAAGCTAAGCGCGAGTTGTTCCCAGACTTTGGCGAAATCG<br>GAACGTTGTGTCGCGATATCTTTTTTCGCAATAATCTTATCATGCGCGCTTGCGGGGATCATATTGTAAGTGCCCCGCCATTGGTGA |

| | |
|---|---|
| | TGACTCGTGCCGAGGTAGATGAGATGTTAGCAGTCGCAGAGCGCTGCCTTGAGGAGTTTGAGCAAACATTAAAAGCTCGCGGACT TGCCTGA |
